# MHC class I Ligands of Rhesus Macaque Killer-Cell Immunoglobulin-Like Receptors

**DOI:** 10.1101/2022.05.25.493479

**Authors:** Jennifer L. Anderson, Kjell Sandstrom, Willow R. Smith, Molly Wetzel, Vadim A. Klenchin, David T. Evans

## Abstract

Definition of MHC class I ligands of rhesus macaque KIRs is fundamental to NK cell biology in this species as an animal model for infectious diseases, reproductive biology, and transplantation. To provide a more complete foundation for studying NK cell responses, rhesus macaque KIRs representing common allotypes of lineage II *KIR* genes were tested for interactions with MHC class I molecules representing diverse *Mamu-A, -B, -E, -F, -I* and *-AG* alleles. KIR-MHC class I interactions were identified by co-incubating reporter cell lines bearing chimeric KIR-CD3ζ receptors with target cells expressing individual MHC class I molecules and were corroborated by staining with KIR IgG-Fc fusion proteins. Ligands for 11 KIRs of previously unknown specificity were identified that fell into two general categories: interactions with multiple Mamu-Bw4 molecules or with Mamu-A-related molecules, including several allotypes of Mamu-AG and the hybrid Mamu-B*045:03 molecule. Although both groups include inhibitory and activating receptors, the majority of KIRs found to interact with Mamu-Bw4 are inhibitory, whereas most of the KIRs that interact with Mamu-AG are activating. We also identified Mamu-A1*012:01 as a ligand for KIR3DLw03*002, which belongs to a phylogenetically distinct group of macaque KIRs with a three amino acid deletion in D0 that is also present in human KIR3DL1/S1 and KIR3DL2. This study more than doubles the number of rhesus macaque KIRs with defined MHC class I ligands and identifies novel interactions with Mamu-AG, -B*045, and -A1*012. These findings support overlapping, but nonredundant, patterns of ligand recognition that reflects extensive functional diversification of these receptors.

## Introduction

Natural killer cell responses in humans and other primate species are regulated in part by interactions between two highly polymorphic sets of molecules; the killer cell Ig-like receptors (KIRs) expressed on the surface of NK cells and their major histocompatibility complex (MHC) class I ligands on target cells (1, 2). As indicated by their nomenclature, KIRs typically have two or three extracellular Ig-like domains (2D or 3D) and either long (L) or short (S) cytoplasmic tails. KIRs with long cytoplasmic tails contain a pair of ITIMs that transduce inhibitory signals, whereas those with short cytoplasmic tails associate with adaptor molecules such as DAP12 or FcεRIγ to transduce activating signals (2, 3). Inhibitory KIRs normally suppress NK cell responses through interactions with MHC class I ligands on the surface of heathy cells. However, if these interactions are somehow perturbed, for instance as a result of MHC class I downregulation in virus-infected cells or deletion of MHC class I genes in tumor cells, the loss of these inhibitory signals can trigger an NK cell response (4–9). Although the molecular mechanisms are not as well defined, activating KIRs can also interact with MHC class I molecules on unhealthy cells to directly trigger an NK cell response (2, 10–12).

Polymorphisms in the *KIR* and *HLA class I* genes have been associated with differences in the course of infection for a number of human viral pathogens and with the outcome of cancer immunotherapy (12–19). *KIR* and *HLA class I* polymorphisms have also been linked to complications during pregnancy and to the success of hematopoietic stem cell and solid organ transplantation (20–23). However, studies to address the immunological mechanisms underlying these observations have been hampered by the lack of a suitable animal model. Mice and other rodents do not have KIRs, but instead express C type lectin-like molecules on their NK cells as polymorphic receptors for MHC class I ligands (2, 24). Moreover, due to the rapid pace of KIR and MHC class I evolution (25–27), it is not possible to predict KIR ligands in non-human primates based on sequence similarity with human KIRs.

In contrast to humans, which have three classical *HLA class I* genes (*HLA-A, -B* and *-C*), Old World monkeys express an expanded array of polymorphic MHC class I genes related to *HLA-A* and *-B*. However, these species do not have an *HLA-C* ortholog, since *MHC-C* arose in hominids as a duplication of an *MHC-B* gene after their divergence from Old World monkeys (28, 29). The *MHC class I* haplotypes of Old World monkeys are polygenic, and rhesus macaques typically express two or three *<u>Ma</u>caca <u>mu</u>latta (Mamu)-A* genes and 4-11 *Mamu-B* genes (30–32). Depending on their transcriptional abundance, the *MHC class I* genes of macaques can be further classified as “major” or “minor” loci (32). Products of the major genes exhibit the greatest polymorphism and present peptides to virus-specific CD8^+^ T cells (33), suggesting functional similarities to HLA-A and -B in humans.

Four non-classical *MHC class I* genes have also been identified in Old World monkeys. Orthologs of *HLA-E* and *-F* (*Mamu-E* and *-F* in rhesus macaques) are well conserved and serve similar functions as their human counterparts (34–37). Macaques and other Old World monkeys also have an ortholog of *HLA-G*; however the *MHC-G* genes of these species have accumulated multiple stop-codons and no longer encode functional proteins (38, 39). In the absence of a functional *MHC-G* gene, Old World monkeys have evolved duplicated *MHC-A* genes, termed *MHC-AG* (*Mamu-AG* in rhesus macaques), that appear to serve a similar function (40–43). Like HLA-G, Mamu-AG has limited polymorphism, a truncated cytoplasmic tail, and is only expressed on placental trophoblasts (40, 41, 44). Mamu-AG is therefore thought to contribute to pregnancy success through interactions with receptors on NK cells and myeloid cells that promote placental vascularization, facilitate fetal tolerance, and protect the maternal-fetal interface from invading microorganisms (44–46). Rhesus macaques also express Mamu-I, which is the product of a fixed duplication of an *MHC-B* gene with limited sequence variation (47).

In concert with the tandem duplication of their *Mamu-A* and *-B* genes, macaques have evolved an expanded complement of lineage II *KIRs* that encode 3D KIRs for Mamu-A and -B ligands (26, 48–52). Whereas humans only have two lineage II *KIR* genes that encode KIR3DL1/S1 and KIR3DL2 as receptors for HLA-Bw4 and –A3/A11, respectively, rhesus macaques have at least 19 lineage II *KIRs* that encode inhibitory and activating receptors for Mamu-A and -B molecules (50, 52–56). Furthermore, in contrast to humans, which have multiple lineage III *KIR* genes encoding KIR2DL/S receptors for HLA-C1 and -C2 ligands (1, 2), macaques only have two lineage III *KIR* genes, one is a pseudogene (KIR3DP1) and the other (KIR1D) expresses a single domain KIR of uncertain function (52). Thus, while human NK cell regulation predominately depends on KIR2DL/S-HLA-C interactions, macaque NK cells are dependent on KIR3DL/S interactions with Mamu-A and -B ligands (52).

The rhesus macaque has become an increasingly valuable animal model for infectious diseases (57–68), reproductive biology (44, 69), and transplantation (70, 71). Although MHC class I ligands have been identified for a few macaque KIRs (72–76), ligands for most of these receptors remain undefined. We therefore performed a systematic survey of MHC class I interactions for common lineage II KIR allotypes representing 16 predicted genes of Indian-origin rhesus macaques. By measuring the responses of KIR-CD3ζ-transduced reporter cell lines to target cells expressing individual MHC class I molecules and confirming interactions by staining with KIR IgG-Fc domain (KIR-Fc) fusion proteins, we identified MHC class I ligands for 11 rhesus macaque KIRs of previously undefined specificity. This represents the most comprehensive analysis of ligand recognition by macaque KIRs to date and reveals novel interactions as well as extensive functional diversification of these receptors.

## Materials and Methods

### Jurkat NFAT luciferase reporter cells expressing chimeric KIR-CD3ζ receptors

cDNA sequences encoding the leader peptide of Mamu-KIR3DL05*008 followed by a Flag-tag (DYKDDDDK) and the D0, D1, D2 and stem domains of rhesus macaque KIRs were synthesized (Integrated DNA Technologies) and cloned into pQCXIP in-frame with sequences encoding the transmembrane and cytoplasmic domains of human CD3ζ. Accuracy of all plasmid constructs was verified by Sanger sequencing of the entire ORF. Retroviral vectors were packaged into VSV G–pseudotyped murine leukemia virus (MLV) particles by co-transfecting GP2-293 cells (Clontech Laboratories) with pQCXIP constructs and pVSV-G (Clontech Laboratories). Jurkat NFAT luciferase (JNL) cells (Signosis) were transduced via spinoculation for one hour with filtered supernatant collected from transfected GP2-293 cells and 2 days later were placed under selection in RPMI medium supplemented with 10% FBS, L-glutamine, penicillin, and streptomycin (R10 medium) plus 0.1 mg/ml hygromycin (Invitrogen) to maintain the luciferase reporter gene. Selection for the KIR-CD3ζ receptor was performed by increasing the puromycin (Invitrogen) concentration to 1.0 µg/ml over 3-4 weeks and then maintaining at 0.5 µg/ml. The GenBank accession numbers for rhesus macaque KIR and human CD3ζ sequences are as follows: *Mamu-KIR3DL02*004:01* (LT963635.1), -*KIR3DLw03*002* (EU419055.1), *-KIR3DL04*001:02* (GU299490.1), *-KIR3DL07*004* (EU419060.1), *-KIR3DL10*002:01* (GU112259.1), *-KIR3DL11*001* (GU112271.1), *-KIR3DLw34*002* (previously -*KIR3DLw03*003*; EU419031.1), -*KIR3DS01*003* (GU564161.1), *-KIR3DS02*004:02* (GU014296.1), -*KIR3DS03*003* (EU702454.1), *-KIR3DS04*003:01* (EU419029.1), -*KIR3DS05*002:01* (GU112262.1), *-KIR3DS06*002:02* (GU112298.1), *-KIR3DSw07*001* (GU112272.1), -*KIR3DSw08*001* (AY505479.1), *-KIR3DSw09*003* (MF164924.1) and *CD3ζ* (J04132.1).

### Rhesus macaque MHC class I-expressing 721.221 cells

Codon-optimized cDNA sequences encoding rhesus macaque MHC class I heavy chain constructs were cloned into pQCXIP or pQCXIN and verified by Sanger sequencing of the entire ORF. Retroviral vectors were packaged into VSV-G-pseudotyped MLV particles as described above. 721.221 cells were transduced via spinoculation for one hour with filtered supernatant collected from transfected GP2-293 cells and placed under selection in R10 medium containing 0.4 mg/ml puromycin or 0.5 mg/ml G418, depending on the vector, to select for MHC class I expression. Selection for the MHC class I receptor was performed for pQCXIN constructs by increasing the G418 (Enzo Life Sciences) concentration to 1.0 mg/ml (based on active units) and then maintaining at 0.5 mg/ml and for pQCXIP constructs by increasing the puromycin (Invitrogen) concentration to 1.0 µg/ml and then maintaining at 0.4 µg/ml. The GenBank accession numbers for rhesus macaque MHC class I sequences are as follows: *Mamu-A1*001:01:01:01* (LT908117), -*A1*002:01:01:01* (LN899621), *-A1*004:01:01:01* (FN396407.1), *-A1*008:01:01:01* (LN899628), *-A1*011:01:01:01* (AJ542579), *-A1*012:01:01:01* (AF157398.1), *-A2*05:02:01:01* (LN899639), *-A2*05:04:01:01* (LT908121.1), *-A3*13:02:01:01* (LN899654.1), *-A3*13:03:01:01* (AF157401), *-A4*14:03:01:01* (GU080236.1), *-A6*01:03:02:01* (EF602318.1), *-B*001:01:02:01* (LM608018), *-B*002:01:01:01* (LN851852), *-B*005:01:01:01* (LM608023), -*B*007:01:01:01* (U41829.1), *-B*008:01:01:01* (U41830), *-B*015:01* (AM902541.1), *-B*017:01:01:01* (AF199358), *-B*022:01:01:01* (LN899675), *-B*036:01:01:01* (AJ556886.1), *-B*041:01:01:01* (LN899682), *-B*043:01:01:01* (AJ556893), *-B*045:03:01:01* (LM608047), *-B*056:01:01:01* (GQ902079.1), *-B*065:01:01:01* (AJ620416.1), *-I*01:01:01* (LT908864), *-E*02:01:02* (previously E*02:01; NM_001114966.1) *-F*01:01* (LT899414), *-AG1*05:01* (FJ409466), *-AG2*01:01* (previously AG*01:01; U84783.1), *AG2*01:02:01:01* (LR989653)*-AG3*02:01:01:01* (U84785.1), *-AG3*03:01:01:01* (U84787.1), -*AG4*08:01:01:01* (LR989670), -*AG4*09:02:02:02* (LR990451), *-AG5*07:06:01:01* (LR989633), *-AG5*07:10:01:02* (LR989642), and *-AG6*04:01:01:01* (LR989697). Only partial-length cDNA sequence was available for *-AG1*05:01.* For the construct expressing this allele, sequences encoding the α3-, transmembrane and cytoplasmic domains were inferred from the *Mamu-AG* consensus sequence.

### KIR-Fc fusion proteins

All KIR-Fc fusion constructs were cloned into an in-house made expression construct (pCIW) that contains the following regulatory elements: CMV promoter/enhancer, SV40 intron, woodchuck hepatitis virus posttranscriptional regulatory element and SV40 polyadenylation signal. The constructs encoded an artificial “secrecon” leader peptide (77) followed by KIR ectodomain fused to the Fc domain of murine IgG2A. Following cloning of the initial KIR3DL01 construct using a synthetic gBlock DNA fragment (IDT DNA), all other KIR fragments were cloned in place of the KIR3DL01 using the “QuickChange cloning” protocol as described by Klenchin et al (78). All fusion proteins were expressed in Expi-293F cells in 250-280 ml culture using the ExpiFectamine 293 Transfection Kit (Life Technologies) according to the manufacturer’s recommendations. Five-six days after transfection, the supernatant was collected, filtered through a 0.45 µM PES filter, and loaded on a rProtein A GraviTrap column (GE Healthcare). After thorough washing of the column with PBS (at least 30 ml), KIR-Fc fusion proteins were eluted with 3 ml Gentle Ag/Ab Buffer (Pierce) and buffer exchanged into 50 mM sodium citrate, (pH 6.5) using three cycles of concentration/dilution in Amicon Ultra-15 50K centrifugal Filter unit (Millipore). The final concentrations of the purified KIR-Fc fusion proteins were calculated from absorbance at 280 nm using theoretical extinction coefficients calculated by ProtParam (https://web.expasy.org/protparam/). After addition of 3 mM sodium azide, the purified protein preparations were clarified by centrifugation at 30,000 *g* for 30 min at 20 °C and the supernatants were snap frozen in liquid nitrogen as 50 µl aliquots and stored at -80 °C.

### Flow cytometry

KIR-CD3ζ expression on the surface of JNL cells and MHC class I expression on 721.221 cells was confirmed by staining with Near-IR fluorescent dye (Invitrogen) followed by 30 min staining at room temperature with PE-conjugated anti-Flag (clone REA216, Miltenyi Biotec) or pan-HLA class I-specific antibodies (clone W6/32; Life Technologies). The cells were then washed twice in PBS plus 1% FBS (FACS buffer) and fixed in 2% formaldehyde in PBS. To assess KIR binding, MHC class I-transduced 721.221 cells were stained for 30 minutes at room temperature with Near-IR fluorescent dye, washed twice with FACS buffer and stained for 30 minutes at room temperature with KIR-Fc fusions proteins (10-20 µg/ml). The cells were then washed twice with FACS buffer, stained for 45 minutes with Alexa Flour 647-conjugated goat anti-mouse IgG (RRID AB_2338925, Jackson ImmunoResearch), washed twice, and stained for 30 minutes at room temperature with the PE-conjugated pan-MHC class I-specific antibody W6/32. The cells were washed again and fixed in 2% formaldehyde. All flow cytometry data were collected using a FACSymphony™ A3 Cell Analyzer (BD Biosciences) and analyzed using FlowJo version 10.8.0. After gating on viable cells, KIR-CD3ζ (Flag-tag) and MHC class I staining was compared to parental JNL and 721.221 cells, respectively. KIR-MHC class I binding was similarly assessed by comparing KIR-Fc versus W6/32 staining on the surface of MHC class I-transduced 721.221 cells to their levels of non-specific staining on parental 721.221 cells.

### KIR-CD3ζ JNL reporter assays

KIR-CD3ζ JNL cells (100, 000) and MHC class I-transduced 721.221 cells (100, 000) were co-cultured overnight at 37 °C and 5% CO2 in R10 medium without antibiotics. KIR-CD3ζ JNL cells were also incubated with parental 721.221 cells as negative control and with anti-Flag-tag (5 µg/ml) (clone 5A8E5, GenScript) and goat anti-mouse (10 µg/ml) (Poly4053, Biolegend) antibodies as a positive “X-link” control. Each combination was tested in triplicate wells (100 µl/well). After 12-18 hours, BriteLite Plus luciferase substrate (PerkinElmer) was added to each well (100 µl/well) and the relative light units (RLU) of luciferase activity was measured using a VICTOR X4 multiplate reader (PerkinElmer).

### Alignments and Phylogenetic trees

Amino acid sequences of KIR and MHC class I molecules were aligned using Geneious Prime 2021.2.2 (Biomatters Ltd). Phylogenetic trees were constructed using raw difference neighbor-joining analysis of nucleic acid sequence for the α1-α3 domains of Mamu-MHC and the D0-D2 domains of human and rhesus macaque KIRs was performed using MAFFT (version 7, publicly available at https://mafft.cbrc.jp/alignment/server/) (79, 80). All analyses utilized G-INS-I iterative refinement with 1000 bootstrap replicates. Trees were viewed using Phylo.io version 1.0.0 (81). For the analysis of KIR sequences, human lineage II KIRs were included as an outgroup (KIR3DL1*001, AY760023.1 and KIR3DL2*001, MN196233.1).

### Statistical analysis

The luciferase induction by KIR-CD3ζ JNL cells (RLU data) was analyzed by one-way ANOVA for multiple comparisons of mean responses to each of the MHC class I-expressing 721.221 cells with mean responses to the parental 721.221 cells. Statistical analyses were performed using GraphPad Prism for Mac OS version 10.8.0.

## Results

Extensive sequencing and segregation analyses of *KIR* transcripts support the existence of at least 22 *KIR* genes in Indian-origin rhesus macaques (53–56). Although our group and others have identified ligands for a few rhesus macaque KIRs (72–76), ligands for most of the KIR in this species remain undefined. To provide a more complete foundation for studying NK cell biology in this species, we investigated the MHC class I interactions of 16 KIRs representing common alleles of distinct lineage II genes in animals of Indian descent (Fig. 1A). Jurkat cells harboring an NFAT-inducible luciferase reporter gene (JNL cells) were transduced with retroviral vectors expressing chimeric KIR-CD3ζ receptors consisting of the extracellular domains of macaque KIRs (D0, D1, D2 and stem domains) fused to the transmembrane and cytoplasmic domains of human CD3ζ. To confirm surface expression, a Flag-tag was appended to the N-terminus of the D0 domain of each KIR-CD3ζ receptor (Fig. 1B). 721.221 cells were similarly transduced with vectors expressing rhesus macaque MHC class I molecules and surface expression was confirmed with a pan-MHC class I antibody (Supplemental Figs 1-3). After co-incubating the KIR-CD3ζ JNL cells overnight with MHC class I-transduced 721.221 cells, ligand recognition was detected by the MHC class I-dependent upregulation of luciferase. KIR-MHC class I interactions were also corroborated by staining MHC class I-expressing 721.221 cells with KIR-Fc fusion proteins consisting of the extracellular domains of macaque KIRs fused to the Fc domain of murine IgG2A.

**FIGURE 1.**
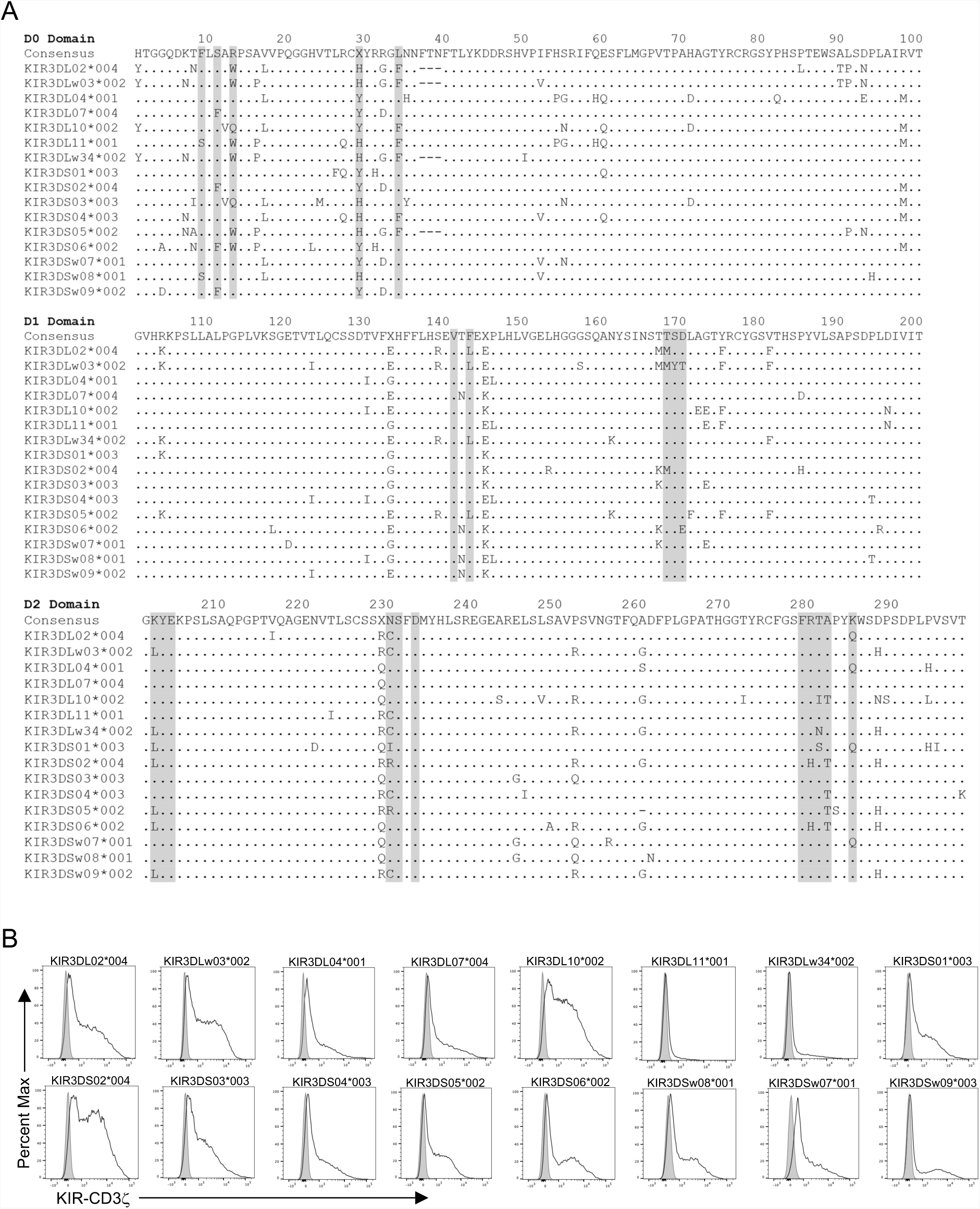
Rhesus macaque KIRs and KIR-CD3ζ expression on JNL cells. (**A**) Alignment of amino acid sequences for the D0, D1 and D2 domains of rhesus macaque KIRs. Shaded sequences correspond to predicted MHC class I contact sites based on the crystal structure of HLA-B*57 in complex with human KIR3DL1 (90). Positions of identity are indicated by periods and amino acid differences are identified by their single-letter code. (**B**) KIR-CD3ζ-transduced JNL cells were stained with anti-Flag antibody and Near-IR LIVE/DEAD stain. After excluding dead cells, the fluorescence intensity of KIR-CD3ζ (Flag-tag) staining on KIR-CD3ζ JNL cells (open) was compared to background staining on parental JNL cells (shaded).

### Mamu-Bw4 recognition by multiple rhesus macaque KIRs

Four of the rhesus macaque KIRs recognized Mamu-Bw4 ligands. KIR3DL04*001- and KIR3DL10*002-CD3ζ JNL cells responded to 721.221 cells expressing Mamu-B*007:01, - B*022:01, -B*041:01, -B*043:01 and -B*065:01 (Fig. 2A, Supplemental Fig. 4A). Similarly, KIR3DL11*001-CD3ζ JNL cells responded to Mamu-B*007:01, -B*041:01 and -B*065:01, and KIR3DS04*003-CD3ζ JNL cells responded to Mamu-B*22:01, -B*041:01 and -B*065:01 (Fig. 2A, Supplemental Fig. 4A). These interactions were corroborated by KIR-Fc staining of 721.221 cells expressing these molecules (Fig. 2B, Supplemental Fig. 4B).

**FIGURE 2.**
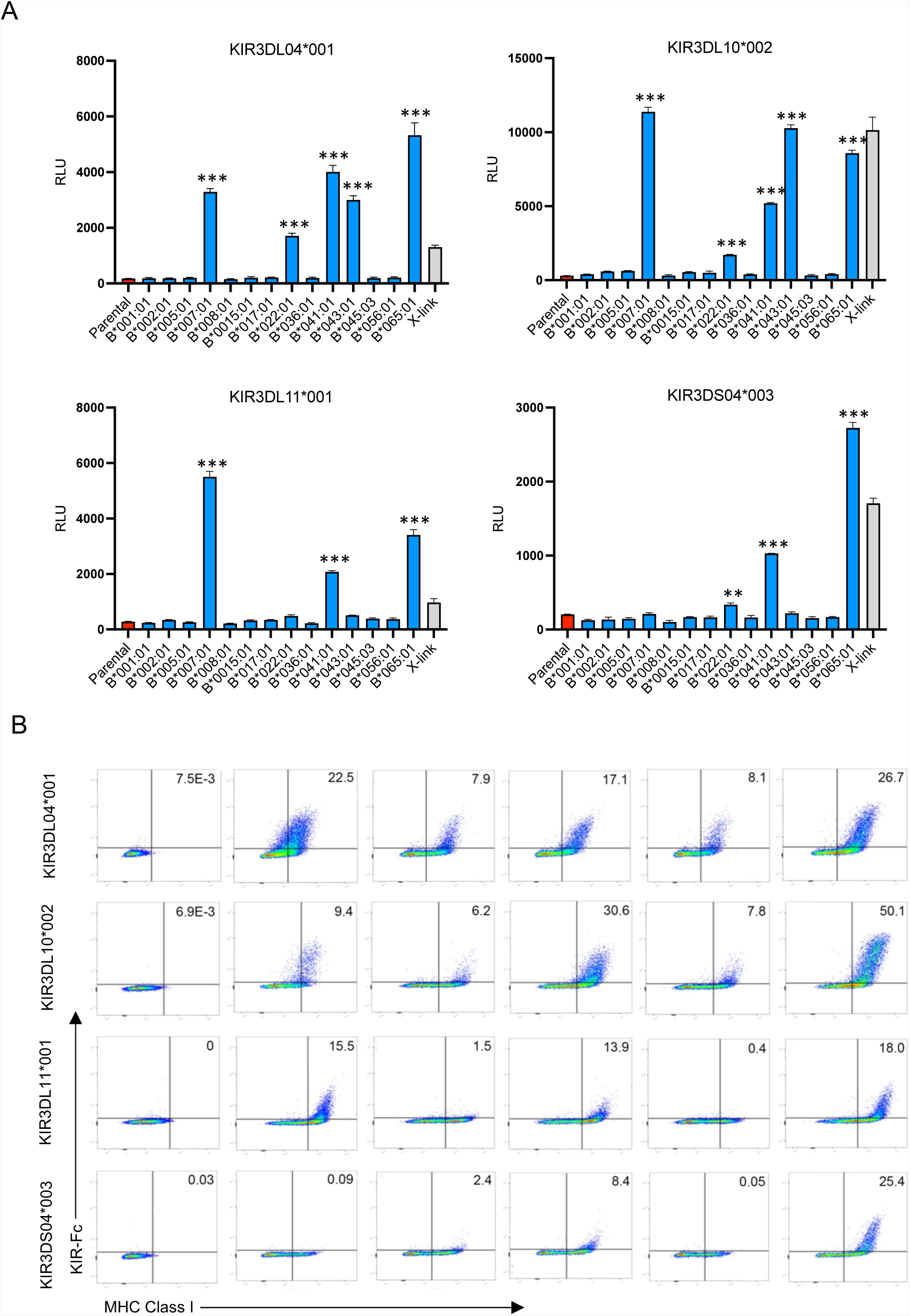
KIR3D04*001, KIR3DL10*002, KIR3DL11*001 and KIR3DS04*003 interact with multiple Mamu-Bw4 ligands. (**A**) KIR3DL04*001-, KIR3DL10*002-, KIR3DL11*001- and KIR3DS04*003-CD3ζ JNL cells were incubated with 721.221 cells expressing the indicated Mamu-B molecules. The bar graphs represent the mean and standard deviation (error bars) of luciferase activity (RLU) from triplicate wells of KIR-CD3ζ JNL cells incubated with Mamu-B + 721.221 cells (blue), parental 721.221 cells (red) and anti-Flag plus anti-mouse antibodies (X-link, gray). Asterisks denote significant differences by one-way ANOVA with Dunnett test (** p=0.0002, and *** p < 0.0001). These results are representative of at least three independent experiments. (**B**) Parental and Mamu-B+721.221 cells were stained with Near-IR LIVE/DEAD dye, KIR3DL04*001-, KIR3DL10*002-, KIR3DL11*001- and KIR3DS04*003-Fc followed by goat anti-mouse IgG and MHC class I-specific antibody (W6/32).

Mamu-B*007:01, -B*022:01, -B*041:01, -B*043:01 and -B*065:01 share a Bw4 motif at residues 77-83 (Supplemental Fig. 2), which was previously shown to contribute to recognition by KIR3DL01 as well as three other rhesus macaque KIRs (73, 75). The identification of Mamu-Bw4 ligands for KIR3DL04*001, KIR3DL10*002, KIR3DL11*001 and KIR3DS04*003 doubles the number of rhesus macaque KIRs known to interact with this group of molecules and reveals an extraordinary expansion of the lineage II KIRs in macaques with this specificity (Table I). Differences in the particular Mamu-Bw4 molecules recognized by these KIRs and the relative strength of these interactions suggests extensive functional diversification of these receptors.

**Table I.** MHC class I ligands of rhesus macaque lineage II KIRs

| KIR Gene<sup>a</sup> | Allotype<sup>b</sup> | MHC class I ligands<sup>c</sup> | Group<sup>d</sup> | Ref<sup>e</sup> |
| --- | --- | --- | --- | --- |
| *KIR3DL01* | KIR3DL01*012 | Mamu-B*007:01, -B*022:01, -B*041:01, -B*043:01, -B*065:01 | Bw4 | (73) |
| *KIR3DL02* | **KIR3DL02\*004** | none detected<sup>f</sup> |  | This study |
| *KIR3DLw03* | **KIR3DLw03\*002** | **Mamu-A1\*012:01** | A | This study |
| *KIR3DL04* | **KIR3DL04\*001** | **Mamu-B\*007:01, -B\*022:01, -B\*041:01, -B\*043:01, -B\*065:01** | Bw4 | This study |
| *KIR3DL05* | KIR3DL05*008 | Mamu-A1*002:01, -A3*13:03, -B*036:01, -AG2*01:01, -AG1*05:01 | AG/A-related | (72, 74, 75) |
| *KIR3DL06* | KIR3DL06*001 | Mamu-B*007:01, -B*022:01, -B*041:01, -B*043:01, -B*065:01 | Bw4 | (75) |
| *KIR3DL07* | **KIR3DL07\*004** | **Mamu-AG2\*01:01, -AG2\*01:02, -AG6\*04:01, -B\*045:03** | AG/A-related | This study |
| *KIR3DL08* | KIR3DL08*002 | Mamu-B*002:01, -B*007:01, -B*022:01, -B*041:01, -B*065:01 | Bw4 | (75) |
| *KIR3DL10* | **KIR3DL10\*002** | **Mamu-B\*007:01, -B\*022:01, -B\*041:01, -B\*043:01, -B\*065:01** | Bw4 | This study |
| *KIR3DL11* | **KIR3DL11\*001** | **Mamu-B\*007:01, -B\*041:01, -B\*065:01** | Bw4 | This study |
| *KIR3DLw34* | **KIR3DLw34\*002** | none detected<sup>f</sup> |  | This study |
| *KIR3DS01* | **KIR3DS01\*003** | **Mamu-AG2\*01:01, -AG6\*04:01** | AG | This study |
| *KIR3DS02* | **KIR3DS02\*004** | **Mamu-AG2\*01:01, -AG2\*01:02,, -AG6\*04:01, -AG1\*05:01, -B\*045:03** | AG/A-related | (76), This study |
| *KIR3DS03* | **KIR3DS03\*003** | none detected<sup>f</sup> |  | This study |
| *KIR3DS04* | **KIR3DS04\*003** | **Mamu-B\*022:01, -B\*041:01, -B\*065:01** | Bw4 | This study |
| *KIR3DS05* | **KIR3DS05\*002** | none detected<sup>f</sup> |  | This study |
| *KIR3DS06* | **KIR3DS06\*002** | **Mamu-AG2\*01:01, -AG1\*05:01, -AG6\*04:01** | AG | This study |
| *KIR3DSw07* | **KIR3DSw07\*001** | **Mamu-AG2\*01:01, -AG6\*04:01** | AG | This study |
| *KIR3DSw08* | **KIR3DSw08\*001** | none detected<sup>f</sup> |  | This study |
| *KIR3DSw09* | **KIR3DSw09\*003** | **Mamu-AG2\*01:01, -AG2\*01:02, -AG1\*05:01, -B\*045:03** | AG/A-related | This study |
| *KIR3DSw39* | KIR3DSw39*002 | Mamu-B*007:01, -B*022:01, -B*041:01, -B*043:01, -B*065:01 | Bw4 | (75) |

<sup>a</sup>Rhesus macaque lineage II KIR genes.
<sup>b</sup>Representative KIR allotypes. KIR allotypes tested in this study are indicated in bold.
<sup>c</sup>Novel MHC class I ligands defined in this study are indicated in bold.
<sup>d</sup>Group of MHC class I ligands.
<sup>e</sup>References for previously reported KIR ligands.
<sup>f</sup>Supplemental Figure 8

### Multiple KIRs recognize Mamu-AG molecules

Six rhesus macaque KIRs were found to interact with Mamu-AG, a non-classical MHC class I molecule expressed at the maternal-fetal interface of the placenta (40–44). JNL cells expressing KIR-CD3ζ chimeras for KIR3DL07*004, KIR3DS01*003, KIR3DS02*004, KIR3DS06*002, KIR3DSw07*001 and KIR3DSw09*003 all responded strongly to 721.221 cells expressing Mamu-AG2*01:01 and -AG6*04:01 (Fig. 3A, Supplemental Fig. 5A). In addition, KIR3DS02*004-CD3ζ JNL cells also engaged 721.221 cells expressing Mamu-AG2*01:02 (Fig. 3A). The binding of these KIRs to Mamu-AG2*01:01 and -AG6*04:01 was corroborated by KIR-Fc staining of Mamu-AG2-expressing 721.221 cells (Fig. 3B, Supplemental Fig. 5B). KIR-Fc staining also revealed detectable interactions with Mamu-AG1*05:01 for KIR3DS02*004, KIR3DS06*002 and KIR3DSw09*003, and Mamu-AG2*01:02 for KIR3DL07*004 and KIR3DSw09*003 that were not observed in cellular assays with KIR-CD3ζ JNL cells (Fig. 3B, Supplemental Fig. 5B). In this case, the detection of binding interactions with Mamu-AG1*05:01 and -AG2*01:02 that were not detectable with KIR-CD3ζ JNL cells may reflect greater sensitivity of KIR-Fc staining or possibly interference of the Flag-tag appended to the N-terminus of the KIR-CD3ζ receptor in the reporter cell assay. These findings identify multiple Mamu-AG ligands for the products of six different KIR genes, five of which encode activating receptors. This brings the total number of rhesus macaque KIRs known to interact with Mamu-AG to seven (Table I).

**FIGURE 3.**
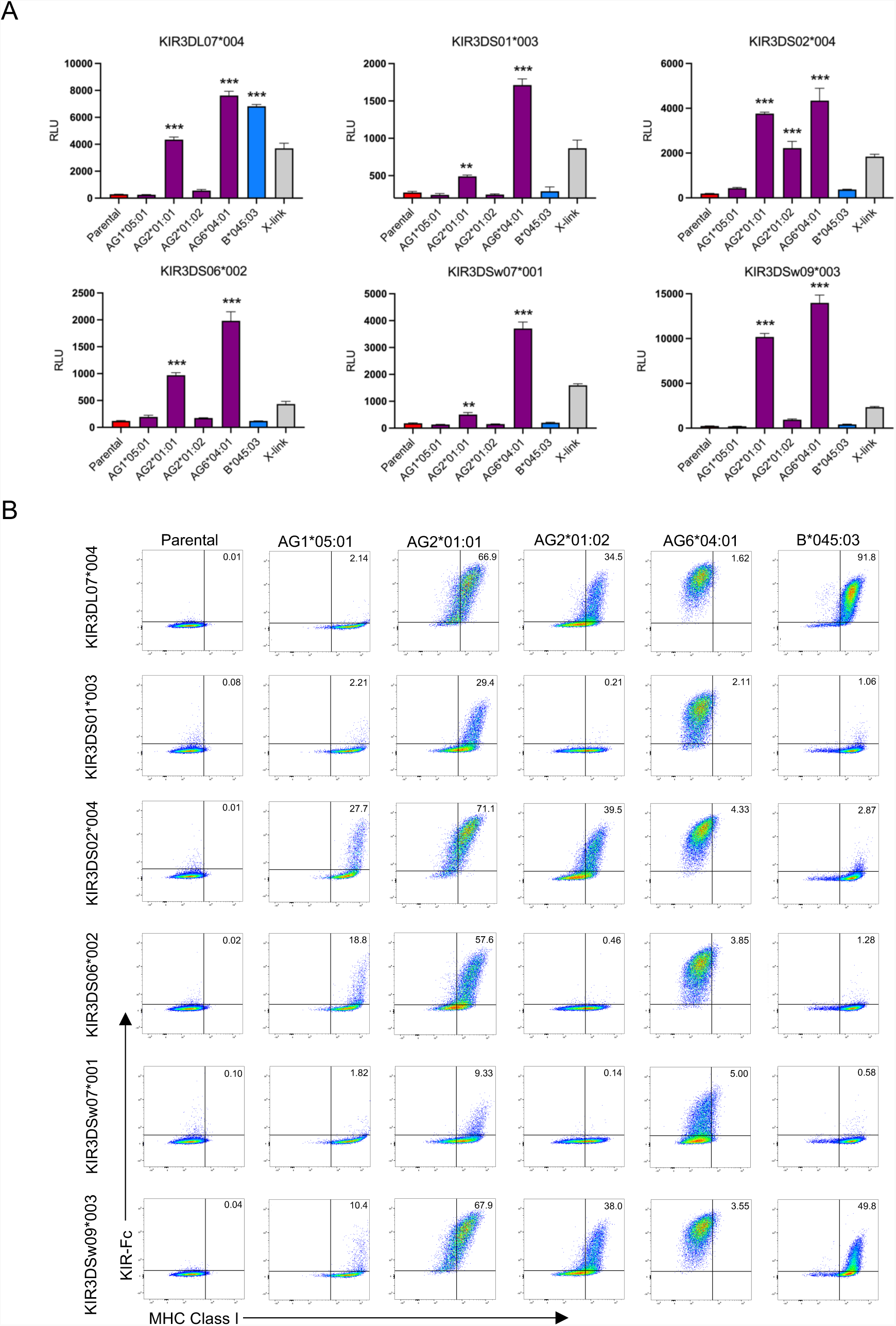
KIR3DL07*004, KIR3DS01*003, KIR3DS02*004, KIR3DS06*002, KIR3DSw07*001 and KIR3DSw09*003 interact with Mamu-AG and -B*045:03. (**A**) KIR3DL07*004-, KIR3DS01*003-, KIR3DS02*004-, KIR3DS06*002-, KIR3DSw07*001- and KIR3DSw09*003-CD3ζ JNL cells were incubated with 721.221 cells expressing the indicated Mamu-AG or -B molecules. The bar graphs represent the mean and standard deviation (error bars) of luciferase activity (RLU) from triplicate wells of KIR-CD3ζ JNL cells incubated with Mamu-AG+721.221 cells (purple), Mamu-B+721.221 (blue), parental 721.221 cells (red) and anti-Flag plus anti-mouse antibodies (X-link, gray). Asterisks denote significant differences by one-way ANOVA with Dunnett test (** p < 0.005, and *** p < 0.0001). These results are representative of at least three independent experiments. (**B**) Parental, Mamu-AG and Mamu-B+721.221 cells were stained with Near-IR LIVE/DEAD dye, KIR3DL07*004-, KIR3DS01*003-, KIR3DS02*004-, KIR3DS06*002-, KIR3DSw07*001- and KIR3DSw09*003-Fc followed by goat anti-mouse IgG and MHC class I-specific antibody (W6/32).

### Recognition of Mamu-B*045:03 by KIR3DL07*004, KIR3DS02*004 and KIR3DSw09*003

In addition to Mamu-AG, JNL cells expressing KIR3DL07*004-CD3ζ also responded to Mamu-B*045:03 (Fig. 3A, Supplemental Fig. 6A). KIR-Fc staining corroborated this interaction and additionally also revealed low but detectable interactions for Mamu-B*045:03 with KIR3DS02*004 and KIR3DSw09*003 (Fig. 3B, Supplemental Fig. 6B). To better understand how these KIR, which predominately recognize Mamu-AG ligands, can also interact with B*045:03, we compared the amino acid sequences for Mamu-B*045:03 with other Mamu-A, -B and -AG molecules. Although Mamu-B*045:03 resembles a typical Mamu-B molecule in its α2 domain, we noted a region of sequence similarity with other Mamu-A molecules in the α1- domain (Fig. 4A). These observations were also supported by phylogenetic analysis. Whereas the full-length nucleotide sequence for Mamu-B*045:03 clusters with Mamu-B molecules (Fig. 4B), the α1-domain is more similar to Mamu-A molecules (Fig. 4C). A similar pattern of clustering was observed for Mamu-B*036, which is a product of recombination between *MHC-A* and *-B* genes and was previously identified as a ligand for KIR3DL05 (75). Mamu-B*045:03 recognition by KIR3DL07*004, KIR3DS02*004 and KIR3DSw09*003 may therefore be explained by its origins as the product of recombination between ancestral *MHC-A* and *-B* genes.

**FIGURE 4.**
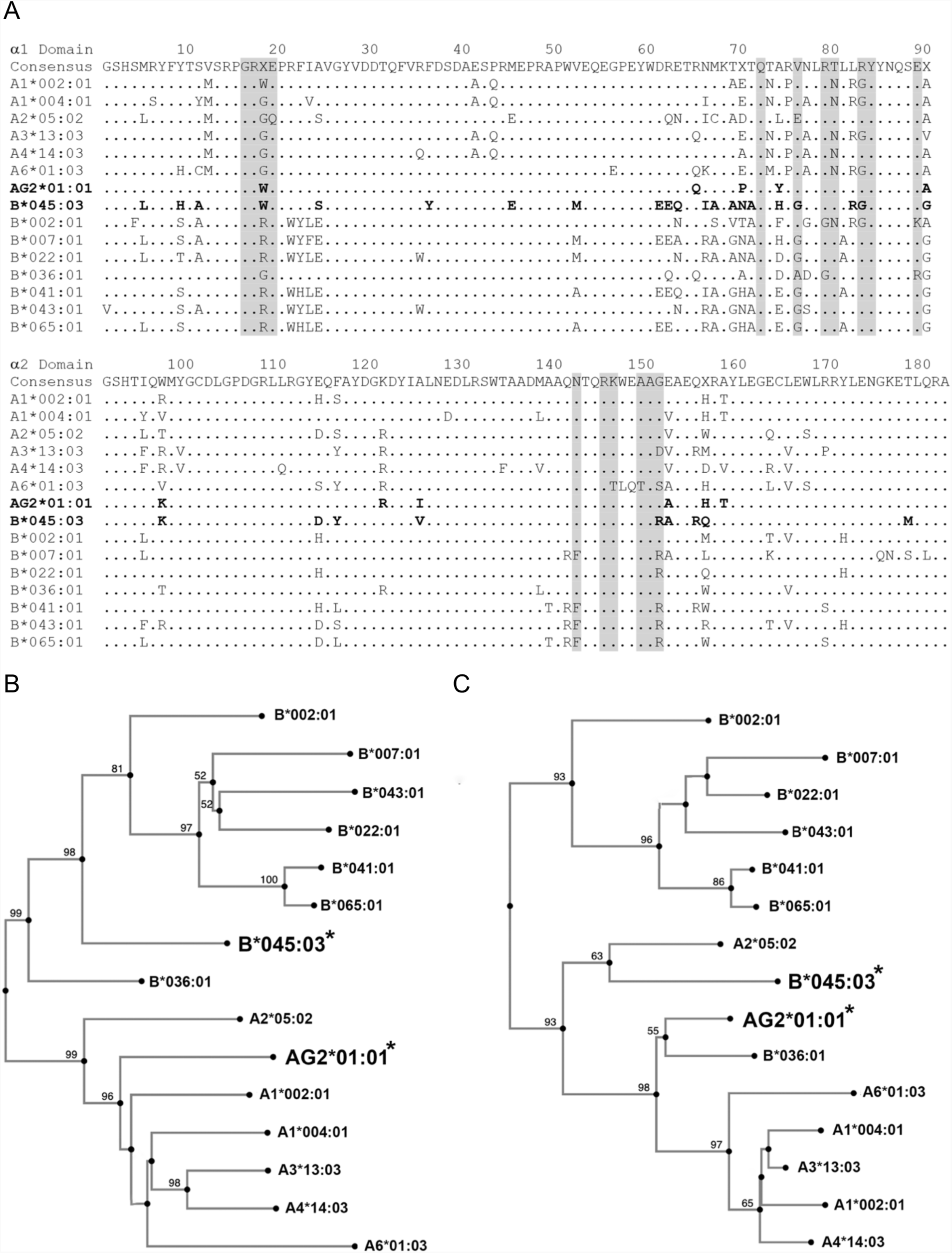
Mamu-B*045:03 has an a1-domain similar to Mamu-A molecules. **(A)** Amino acid sequence alignment for the α1- and α2-domains of rhesus macaque MHC class I molecules. Residues 77–83 corresponding to the Bw4/Bw6 motif are underlined and predicted KIR contact sites based on the crystal structure of HLA-B*57 in complex with human KIR3DL1 are shaded (90). Positions of identity are indicated by periods and amino acid differences are identified by their single-letter code. **(B & C)** Phylogenetic trees of nucleotide sequence coding for the α1-α3 domains **(B)** or the α1 domain (**C**) of selected Mamu-A, -B and -AG molecules. Neighbor-joining analysis was performed using MAFFT software (version 7, publicly available at https://mafft.cbrc.jp/alignment/server/). Bootstrap support of greater than 50% is indicated.

### Recognition of Mamu A1*012:01 by KIR3DLw03*002

KIR3DLw03*002-CD3ζ JNL cells robustly responded to 721.221 cells expressing Mamu- A*012:01 (Fig. 5A, Supplemental Fig. 7A), but not to other MHC class I molecules tested in this study. The binding of KIR3DLw03*002 to Mamu-A*012:01 was also corroborated by KIR-Fc staining (Fig. 5B, Supplemental Fig. 7B). KIR3DLw03*002 is one of four KIRs that display an absence of 3 amino acids at positions 37-39 of the D0 domain compared to the other rhesus macaque KIRs tested in this study (Fig. 1A).

**FIGURE 5.**
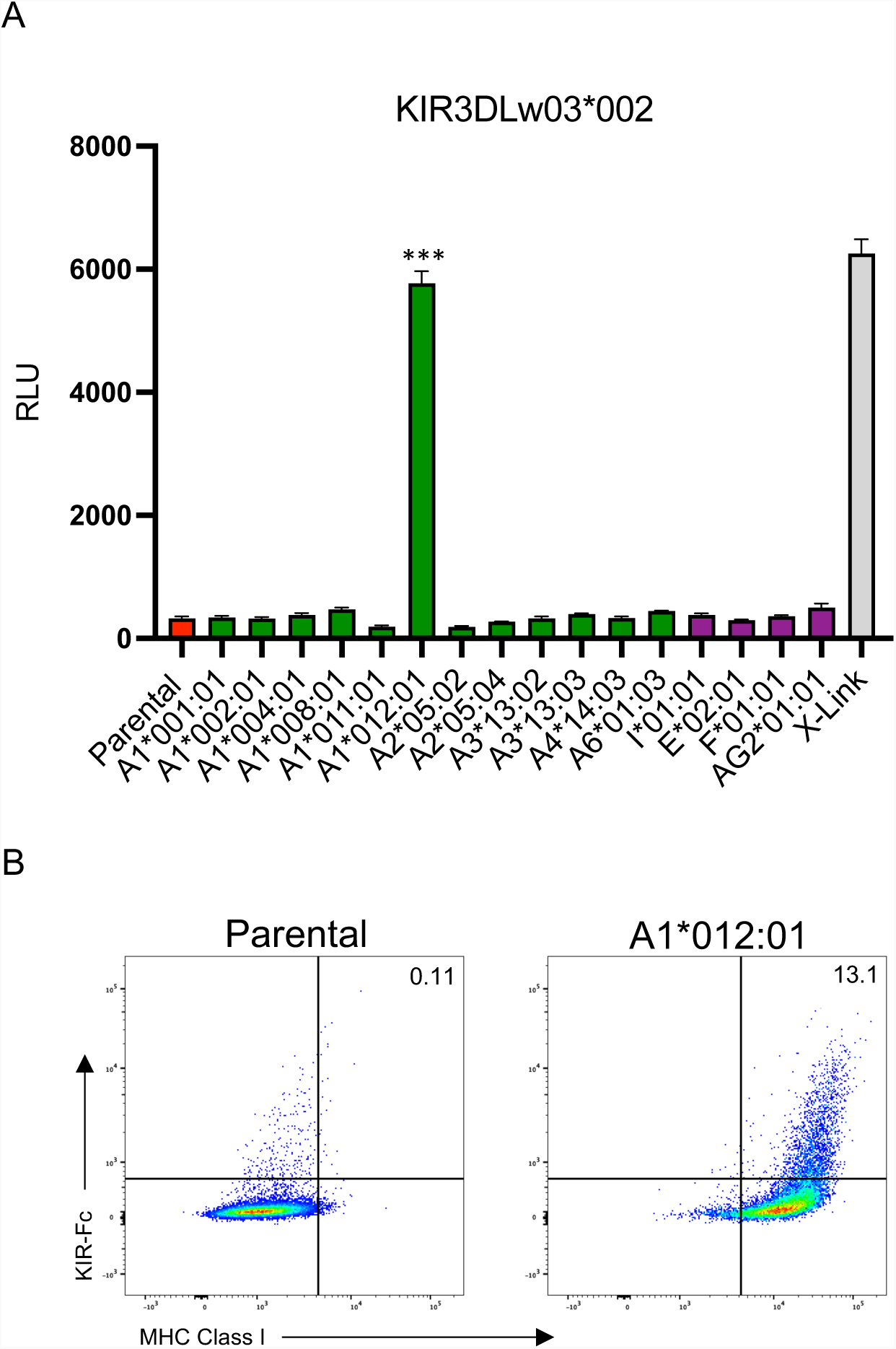
KIR3DLw03*002 interacts with Mamu-A*012:01. (**A**) KIR3DLw03*002-CD3ζ JNL cells were incubated with 721.221 cells expressing the indicated rhesus macaque MHC class I molecules. The bar graphs represent the mean and standard deviation (error bars) of luciferase activity (RLU) from triplicate wells of KIR-CD3ζ JNL cells incubated with Mamu- A*012:01+721.221 (green), nonclassical (purple), parental 721.221 cells (red) and anti-Flag plus anti-mouse antibodies (X-link, gray). Asterisks denote significant differences by one-way ANOVA with Dunnett test (*** p < 0.0001). These results are representative of at least three independent experiments. (**B**) Parental and Mamu-A1*012:01+721.221 cells were stained with Near-IR LIVE/DEAD dye, and KIR3DLw03*002-Fc followed by goat anti-mouse IgG and MHC class I-specific antibody (W6/32).

### Phylogenetic relationship of rhesus macaque KIRs is concordant with patterns of ligand recognition

To investigate the relationship between the extracellular domains of rhesus macaque KIRs and their patterns of ligand recognition, we constructed a phylogenic tree of the D0-D2 domains of rhesus macaque KIRs using human KIR3DL1 and KIR3DL2 as outgroups. The clustering of macaque KIRs generally reflected their ligand recognition. Six of the eight KIRs found to interact with Mamu-Bw4 ligands clustered together on a branch that includes KIR3DSw39*002, KIR3DL06*001, KIR3DS04*003, KIR3DL11*001, KIR3DL04*001 and KIR3DL08*002 (Fig. 6). Likewise, six of the seven KIRs found to interact with Mamu-AG ligands clustered together on a separate branch that includes KIR3DS06*002, KIR3DS01*003, KIR3DL07*004, KIR3DSw09*003, KIR3DS02*004 and KIR3DL05*008 (Fig. 6). KIR3DL10*002, KIR3DSw07*001 and KIR3DLw03*002 were exceptions. Although KIR3DL10*002 and KIR3DSw07*001 interact with Mamu-Bw4 and -AG ligands, respectively, these receptors grouped together on the same branch (Fig. 6). In contrast, KIR3DLw03*002, which interacts with Mamu-A1*012:01, was found on a distinct branch with three other KIRs for which ligands have not yet been identified (KIR3DS05*002, KIR3DLw34*002 and KIR3DL02*004) (Fig. 6). This group of KIRs is separated from the others by strong bootstrap support and is distinguished by the absence of three amino acids (residues 37-39) in the D0 domain (Fig. 1A) that are also missing in human KIR3DL1/S1 and KIR3DL2.

**FIGURE 6.**
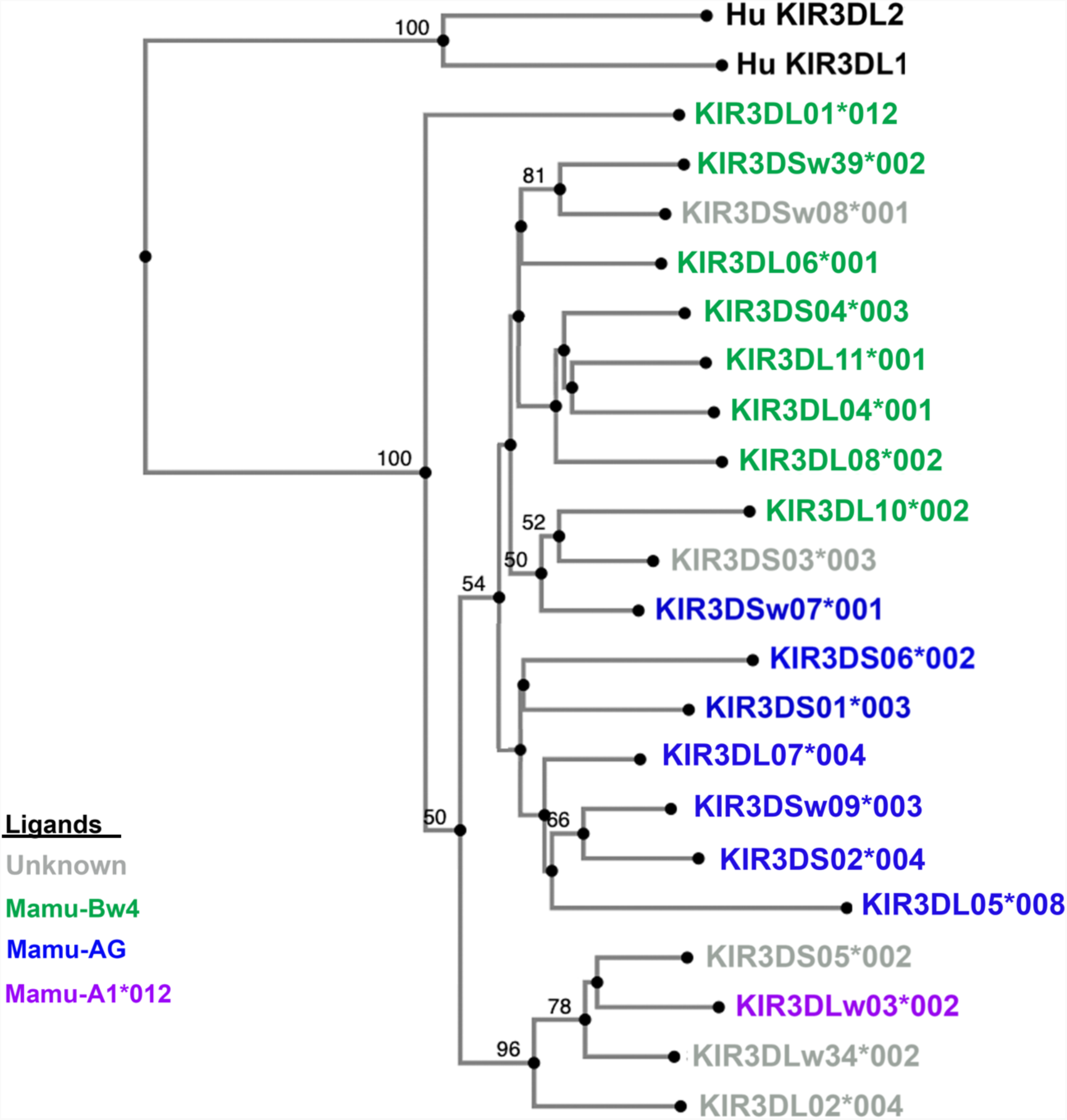
Phylogenetic clustering of rhesus macaque KIRs reflect their patterns of ligand recognition. Neighbor-joining analysis of nucleotide sequences coding for the D0-D2 domains of human and rhesus macaque KIRs was performed using MAFFT software. Bootstrap support of greater than 50% is indicated. Human KIR3DL1 and KIR3DL02 were used included as an outgroup. Rhesus macaque KIRs are color coded according to their MHC class I ligands: unknown (gray), Mamu-Bw4 (green), Mamu-AG (dark blue), and Mamu-A1*012:01 (purple).

## Discussion

Establishing the MHC class I ligands of rhesus macaques KIRs is fundamental to NK cell biology in this species as a preclinical model for infectious diseases, transplantation, and reproductive biology. Because of the rapid pace of KIR evolution in primates, it is not possible to predict the ligands for macaque KIRs on the basis of sequence similarity with their human counterparts. The *KIR* genes of macaques are evolving at a particularly rapid pace characterized by extensive gene duplication and recombination, generating allelic variation and haplotype diversity that greatly exceeds the genetic diversity of human *KIRs* (55, 56, 82). Although the number of alleles and predicted *KIR* genes varies among geographically distinct populations (83), extensive sequencing and segregation analyses strongly support the existence of at least 22 distinct *KIR* genes in rhesus macaques of Indian origin (55, 56, 82). Of these, 19 are predicted to encode lineage II KIRs for Mamu-A and -B ligands. To provide a foundation for studying NK cell responses in macaques, we selected common allotypes representing 16 lineage II KIRs for analysis of their MHC class I interactions. By screening 721.221 cells expressing MHC class I molecules corresponding to 39 *Mamu-A, -B, -E, -F, -I* and *-AG* alleles in assays with KIR-CD3ζ JNL reporter cell lines and by staining these MHC class I-expressing 721.221 cells with KIR-Fc fusion proteins, we identified ligands for 11 previously uncharacterized KIRs.

These analyses revealed two general patterns of ligand recognition; interactions with Mamu-Bw4 molecules and interactions with Mamu-A-related molecules, particularly Mamu-AG. Of the 16 KIRs tested in this study, four (KIR3DL04*001, KIR3DL10*002, KIR3DL11*001 and KIR3DS04*003) were found to interact with molecules that contain a Bw4 motif at residues 77-83. This demonstrates a remarkable expansion of Bw4 specificity in the rhesus macaque. In contrast to humans, which only have a single *KIR3DL1/S1* gene that encodes receptors for HLA-Bw4, rhesus macaques have at least eight lineage II *KIRs* that encode receptors for Bw4 molecules (Table I). Six of the KIRs in this study (KIR3DL07*004, KIR3DS01*003, KIR3DS02*004, KIR3DS06*002, KIR3DSw07*001 and KIR3DSw09*003) were found to interact with Mamu-AG2*01:01 and AG6*04:01, either alone or in combination with Mamu-AG1*05:01, -AG2*01:02, and -B*45:03. Together with earlier studies defining similar ligands for KIR3DL05 (72, 74, 76), at least eight rhesus macaque *KIRs* are known to encode receptors for Mamu-A or -AG ligands (Table I). We also identified Mamu-A1*012:01 as a ligand for KIR3DLw03*002, which belongs to a distinct group of receptors with a characteristic deletion of three amino acids in the D0 domain.

All the KIRs that interact with Bw4 ligands recognize Mamu-B*041:01 and -B*065:01 with varying efficiency, suggesting that these molecules may provide a common reference for this group of receptors (Table I). However, interactions with other Mamu-Bw4 molecules were more variable. Several KIRs, including KIR3DL11*001 and KIR3DS04*003, lack detectable responses to Mamu-B*007:01, -B*022:01 or -B*043:01. Differences in the magnitude of KIR-CD3ζ JNL cell responses and KIR-Fc staining also suggest variation in the strength of these interactions. Although most of the KIRs responding to Mamu-Bw4 ligands are inhibitory, interactions with a couple of activating KIRs (KIR3DS04*003 and KIR3DSw39*002) were also observed (Table I). Thus, differences in the preferential recognition of individual Mamu-Bw4 molecules, variation in the strength of these interactions, and the identification of inhibitory and activating KIRs that respond to this group of molecules, indicates extensive functional diversification of this group of receptors.

Rhesus macaque MHC class I haplotypes typically express two or three *Mamu-A* genes and 4-11 *Mamu-B* genes that can be classified as major or minor loci based on their transcriptional abundance (30–32). The Mamu-Bw4 molecules identified as KIR ligands are products of transcriptionally abundant genes (32). However, phylogenetic and segregation analyses suggest that they are distinct from the *Mamu-B* genes that encode molecules known to present viral peptides to CD8^+^ T cells (30–33, 84). This is further supported by unique polymorphisms in the cytoplasmic domains of Mamu-B*007:01, -B*022:01, -B*041:01, - B*043:01 and -B*065:01 that are not present in the cytoplasmic tails of molecules such as Mamu-B*008 and -B*017 that are known to present SIV epitopes for recognition by CD8+ T cells (84, 85). We previously demonstrated that these cytoplasmic domain polymorphisms prevent MHC class I downmodulation by SIV Nef (86). Selective downmodulation of HLA-A and -B (but not HLA-C) by HIV-1 Nef reduces the susceptibility of virus-infected cells to CD8+ T cells (6, 7, 87). Although the Vpu proteins of certain HIV-1 isolates may downmodulate HLA-C at later stages of infection (88, 89), HLA-C is initially retained on the cell surface, presumably to maintain interactions with inhibitory KIR (6). In a similar manner, SIV Nef downmodulates Mamu-A and -B molecules known to present viral peptides to CD8+ T cells, but not the products of *Mamu-Bw4* genes that serve as ligands for multiple KIRs (75, 86). Hence, parallels in both immunogenetics and virology suggest functional specialization, whereby the products of certain *Mamu-B* genes play a dominant role in restricting CD8+ T cell responses while others serve as KIR ligands to modulate NK cell responses.

Another group of rhesus macaque KIRs interacts with Mamu-A-related ligands, including certain Mamu-AG allotypes and in some cases Mamu-B*045:03. KIR3DL07*004, KIR3DS01*003, KIR3DS02*004, KIR3DS06*002, KIR3DSw07*001 and KIR3DSw09*003 all recognize Mamu-AG2*01:01 and -AG6*04:01 (Table I). Additional interactions with Mamu-AG1*05:01 were detectable for KIR3DS02*004, KIR3DS06*002 and KIR3DSw09*003 and with Mamu-AG2*01:02 for KIR3DL07*004, KIR3DS02*004, and KIR3DSw09*003 (Table I). Mamu-B*045:03 was also identified a ligand for KIR3DL07*004, KIR3DS02*004 and KIR3DSw09*003 (Table I). Although Mamu-B*045:03 is a product of a *Mamu-B* allele, a region of its α1-domain appears to be derived from an *MHC-A* gene. Thus, the hybrid origins of Mamu-B*045:03 probably account for its recognition by these KIRs.

In contrast to KIRs with specificity for Mamu-Bw4, most receptors for Mamu-AG are activating. Five of the seven rhesus macaque KIRs that interact with Mamu-AG2*01:01 and - AG6*04:01 are products of *KIR3DS* genes (Table I). In humans, the ligation of activating lineage III KIRs (KIR2DS) and KIR2DL4 on maternal NK cells of the placental decidua by HLA-C and soluble HLA-G stimulates the release of pro-inflammatory and pro-angiogenic factors that promote placental vascularization (20, 21, 45). Since macaques do not express orthologs of HLA-C or -G (28, 29, 38), it is conceivable that KIR3DS interactions with Mamu-AG play a similar role in placental development. Unlike human *KIRs*, which can be broadly categorized into inhibitory or activating haplotypes based on fixed differences in the number of activating genes, *KIR* haplotypes in macaques are much more variable (82). There are typically 4-17 expressed *KIR* genes per haplotype in the rhesus macaque, including at least one activating gene (56, 82). The expansion of lineage II *KIR* that encode receptors for Mamu-AG may therefore ensure that every animal has at least one activating KIR capable of engaging this non-classical molecule in the placenta.

The identification of Mamu-A1*012:01 as a ligand for KIR3DLw03*002 provides at least one MHC class I ligand for a phylogenetically distinct group of receptors with a characteristic deletion of residues 37-39 in D0. Additional comparison of the D0 domains of KIRs expressed in other primate species, revealed that these residues are also absent from the lineage II KIRs of apes and humans. This suggests that hominid lineage II KIRs evolved from a *KIR* gene with a corresponding 9 base pair deletion in exon 3 that was present in the common ancestor of apes and Old World monkeys. Further inspection indicates that this deletion results in polymorphic differences in potential N-linked glycosylation (PNG) sites that are close to a predicted MHC class I contact site (90). Whereas most rhesus macaque KIRs have two tandem PNG sites (<u>N</u>_36_FT<u>N</u>_39_FT), KIR3DL02, KIR3DLw03, KIR3DLw34 and KIR3DS05 only contain a single PNG site in this region (<u>N</u>_36_FT) (Fig. 1). Furthermore, because of an additional threonine-to-methionine substitution at residue 38 (N_36_FM), human lineage II KIRs (KIR3DL1 and KIR3DL2) lack a PNG site in this region. Hence, these species-specific differences in glycosylation may influence MHC class I interactions with lineage II KIRs.

The present study more than doubles the number of rhesus macaque KIRs with defined MHC class I ligands. At least one MHC class I ligand has now been identified for KIRs representing 16 different lineage II *KIR* genes in animals of Indian origin. These findings reveal a remarkable expansion of lineage II KIRs with broad specificity for Mamu-Bw4 or -AG ligands and novel receptor interactions with Mamu-AG, -B*045 and -A1*012 allotypes. The recognition of overlapping, but non-redundant sets of ligands by both inhibitory and activating KIRs indicates extensive diversification of these receptors in concert with an expanded complement of *Mamu-A* and *-B* genes. These observations advance our basic understanding of KIR and MHC class I co-evolution in primates and provide a valuable foundation for studying NK cell responses in the rhesus macaque as a pre-clinical model.

## Footnotes

This work was supported by Public Health Service grants AI095098, AI155163, AI161816, AI121135, AI148379 and AI098485 to DTE. Additional support was provided by PHS grant OD011106 to the WNPRC. DTE is an Elizabeth Glaser Scientist of the Elizabeth Glaser Pediatric AIDS Foundation and The UW Medical Foundation Professor of Pathology and Laboratory Medicine.

## Abbreviations used in this article

KIR: killer-cell Ig-like receptor
MHC: major histocompatibility complex
*Mamu*: *Macaca mulatta*
JNL: Jurkat NFAT luciferase

