## Supplemental Files for "MHC class I Ligands of Rhesus Macaque Killer-Cell Immunoglobulin-Like Receptors"

Supplemental Figure 1

**A**

```
α1 Domain         10        20        30        40        50        60        70        80        90
Consensus GSHSMRYFYTSMSRPGRGEPRFIAVGYVDDTQFVRFDSDAASQRMEPRAPWVEQEGPEYWDRETRNMKTETQNAPVXLRNLRGYYNQSEA
A1*001:01 .....K............Q.........................................................N..T.LR.......
A1*002:01 .................W..................................................A.......N.............
A1*004:01 .....S....Y...........V..........................................I..........A.............
A1*008:01 ....L.....AV......Q....S................E.P.E.................N..IY.AA...YREG.Q...........
A1*011:01 ........H.AV..........FT..................P.K....................IS.AN..TYREN..TALR.......
A1*012:01 ..................Q..................................................A..TYRES.............
A2*05:02  ....L.............Q....S................E.P.E................QN..IC.AD..TLREN..T.LR.......
A2*05:04  ....L.............Q....S................E.P.E......M.........QN..IC.AD..TLREN..T.LR.......
A3*13:02  .....................................................................A......A............V
A3*13:03  ................................................................Q...........A............V
A4*14:03  ..................................Q..................................A......A....LR.......
A6*01:03  ........H.C.............................E.P............E........QK......M...A.............

α2 Domain        100       110       120       130       140       150       160       170       180
Consensus GSHTFQRMYGCDLGPDGRLLRGYEQYAYDGRDYIALNEDLRSWTAADMAAQNTQRKWEAAGVAEQHRAYLEGECVEWLRRYLENGKETLQRA
A1*001:01 ....L...V.....................K................V............D...SM......Q.....P....K.......T
A1*002:01 ....I..................H.S....K..............................E.....T......L.................
A1*004:01 ....Y.V..................F....K......D.........L...................T......L.................
A1*008:01 ....Y.T................D.S...................................E..RF.T........................
A1*011:01 ......T................D.....................G.....................T......L.................
A1*012:01 .......................S.............G.......................E..RF.T.VD.R...................
A2*05:02  ....L.T................D.S.......................................W......Q.L.S...............
A2*05:04  ....I.T................D.S.......................................W........L.....H...........
A3*13:02  ........V......................................V............D...RM......R...S...............
A3*13:03  ........V...................................................D...RM............P.............
A4*14:03  ........V..........Q.....F.................F...V.................D.V....R...................
A6*01:03  ....I.V................S...............................TLQT.SA..........L...S...............

α3 Domain      190       200       210       220       230       240       250       260       270
Consensus DPPKTHVTHHPVSDHEATLRCWALGFYPAEITLTWQRDGEDQTQDTELVETRPAGDGTFQKWAAVVVPSGKEQRYTCHVQHEGLPEPLTLRW
A1*001:01 ......................................................................E..............K.H..K.
A1*002:01 ..............Q.....................................................................R.......
A1*004:01 ..............Y.........................................................H...................
A1*008:01 ............................................................................................
A1*011:01 ...................................R........................................................
A1*012:01 ............................................................................................
A2*05:02  .S.................................R........................................................
A2*05:04  .S.................................R........................................................
A3*13:02  ........................................E.............................E..S...........K......
A3*13:03  ........................................E.............................E..S...........K......
A4*14:03  .....................................................................................K......
A6*01:03  ...........I.........................................................................K......
```

**B**

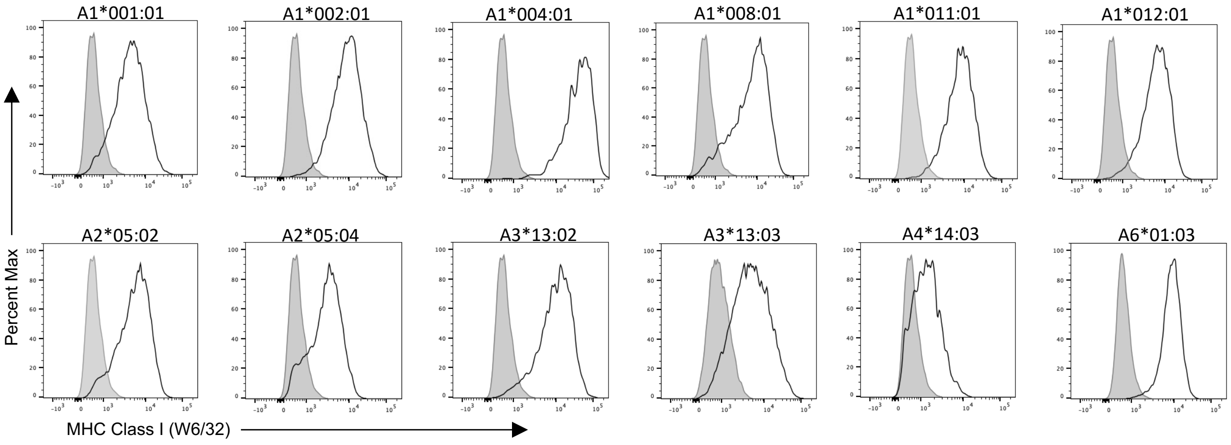


**Supplemental Figure 1.** (A) Amino acid alignment of the α1- α3 domains of Mamu-A molecules used in this study. Residues 77–83 corresponding to the Bw4/Bw6 motif are underlined and predicted KIR contact sites based on the crystal structure of HLA-B*57 in complex with human KIR3DL1 are shaded (90). (B) Surface expression of MHC class I molecules on 721.221 cells. 721.221 cell lines expressing the indicated rhesus macaque MHC class I molecules (open) were stained with a PE-conjugated pan-MHC class I–specific mAb (clone W6/32) and compared with staining on parental 721.221 cell lines (shaded).

Supplemental Figure 2

**A**

```
α1 Domain          10        20        30        40        50        60        70        80        90
Consensus GSHSMRYFSTAVSRPGRREPRFIEVGYVDDTQFVRFDSDAESPRMEPRAPWMEQEGPEYWEEQTRRAKGXAQTDRGNLRTLLRYYNQSEG
B*001:01  ........H........G.....S..................R........I........DRE..K...N.....E...IA.S......T
B*002:01  ...F................WYL............................V........DRN..NS.VT...F.VG.GN.RG.....KA
B*005:01  ....LG..Y........G..L..A......................A....I..............V..H...F.VS..N.RG.....KA
B*007:01  ....L...............WYF.......................................A......N...H......A.........
B*008:01  ..........S......G.....S....................E.................E......H.....AD.G..RG......A
B*015:01  ........T.VM...D...AWYL................V...........V.............NS..H...E.E..............
B*017:01  .....K..Y.S......G.....S....................E......V..........A.....EA...H.E....A........A
B*022:01  ....L...T...........WYL...........W.........................DRN.....AN..........A.........
B*036:01  ........Y.S......G.....A...........................V........DQE.QNM.TAT....AD.G.........R.
B*041:01  ..........S.........WHL............................A.............I...H...E......A.........
B*043:01  V...................WYL...........W................V........DRN......N...E..S.............
B*045:03  ....L...H........W.....S...........Y........E....................I..AN...H.......RG.......
B*056:01  .........N.......G.....A...............V..........................V.EA...F......A........A
B*065:01  ....L.....S.........WHL............................A..........E......H...E......A.........

α2 Domain         100       110       120       130       140       150       160       170       180
Consensus GSHTLQWMYGCDLGPDGRLLRGYXQYAYDGKDYIALNEDLRSWTAADMAAQNTQRKWEAAREAEQLRAYLEGECVEWLRRYLENGKETLQRA
B*001:01  ......M.H..............Y.R..............H......L............GV...R......R..........K........
B*002:01  .......................H.F..................................G....M......T.......H...........
B*005:01  ......R.S..YVE.........D......R................L.............V............P......M..Q....P..
B*007:01  .......................E.F........................RF.........A..........K.L.........QN.S.L..
B*008:01  ......T................H.........F.............V...I.............V......T...................
B*015:01  ....I..................D.S...........Q.....I................GV..........R...................
B*017:01  ....I.K................H.S...........G....................GN.Y..RF........L.................
B*022:01  ....I..................H.F.......................................Q........L.....H...........
B*036:01  ....I.T................E.F....R................L............G....W..........................
B*041:01  ....I..................H.L......................T.RF............RW........L...S.............
B*043:01  ....F.R................D.S........................RF.............R......T.......H...........
B*045:03  ....I.K................D..........V..........................A..RQ........L............M....
B*056:01  ....F...F.....S........E................S......V............G...RM.....................M....
B*065:01  .......................D.L......................T.RF.............W........L...S.............

α3 Domain       190       200       210       220       230       240       250       260       270
Consensus DPPKTHVTHHPVSDHEATLRCWALGFYPAEITLTWQRDGEDQTQDTELVETRPGGDGTFQKWGAVVVPSGEEQRYTCHVQHEGLPEPLTLRW
B*001:01  ................TI..........................................................................
B*002:01  ......................V..............................A......................................
B*005:01  .T.........I............................E........................................K..T..R....
B*007:01  ..............................................................A.............................
B*008:01  ............................................................................................
B*015:01  ........................................E............A..................H...................
B*017:01  ........................................E......F............................................
B*022:01  ...........I....S............................................................Y..............
B*036:01  ...........I............................................................H...................
B*041:01  .....Y......................................................................................
B*043:01  ........................................E...................................................
B*045:03  ........................................E.....................A........................V....
B*056:01  .................................................................................Q..........
B*065:01  .....Y......................................................................................
```

**B**

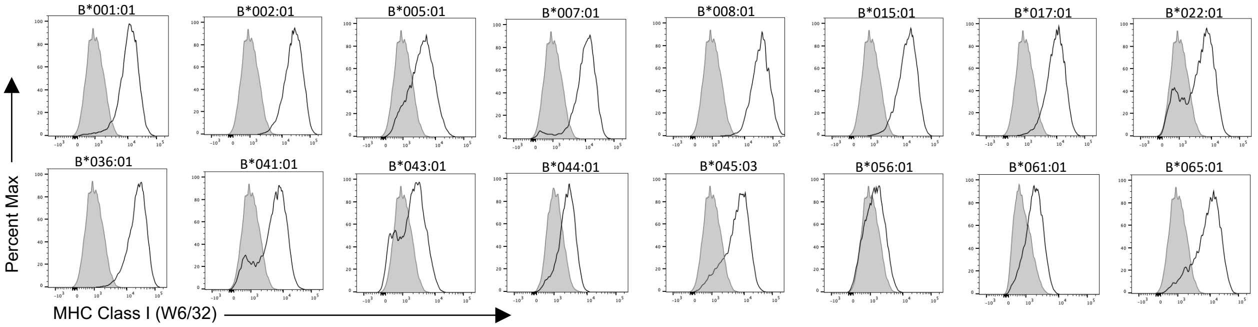


**Supplemental Figure 2.** (A) Amino acid alignment of the α1- α3 domains of Mamu-B molecules used in this study. Residues 77–83 corresponding to the Bw4/Bw6 motif are underlined and predicted KIR contact sites based on the crystal structure of HLA-B*57 in complex with KIR3DL1 are shaded (90). (B) Surface expression of MHC class I molecules on 721.221 cells. 721.221 cell lines expressing the indicated rhesus macaque MHC class I molecules (open) were stained with a PE-conjugated pan-MHC class I–specific mAb (clone W6/32) and compared with staining on parental 721.221 cell lines (shaded).

Supplemental Figure 3

**A**

```
α1 Domain          10        20        30        40        50        60        70        80        90
Consensus  GSHSMRYFYTSVSRPGRGEP--RFIAVGYVDDTQFVRFDSDAESPRMEPRAPWVEQEGPEYWDRETQNMKTATQTYRVNLRTLLRYYNQSEA
AG1*05:01  ..........A......R..--L...........................................................A........G
AG2*01:01  .................W..--.................................................P....................
AG2*01:02  ..........A......W..--.......................................................E..............
AG3*02:01  ...........M......Q.--.......................................................E..............
AG3*03:01  ..........A.......Q.--......................................................QA..............
AG4*08:01  ...........M......Q.--................................................N......G..............
AG4*09:02  ....L...............--..F.............................................N......E..............
AG5*07:06  ....L......M.....R..--..............................................................P.......
AG5*07:10  ....L......M.....R..--L...................G.................................................
AG6*04:01  ..........A......W..--..............Q.......T...............................................
E*02:01    ....LK..H.........G.--...S...........Y....A....................Q..RSARDTA..F....E..RG.......
F*02:01    ....L...S.A......R..RY.Y...E.......L......AI...............Q..E.T.GYA.ANAR.D..A..K..LR......
I*01:01    ....L...G.T.........--...............................M........EE..RIA.ARA..E.G....A.........

α2 Domain          100       110       120       130       140       150       160       170       180
Consensus  GSHTXQKMYGCDLGPDGRLLRGYEQFAYDGRDYIALNEDLRSWTAADMAAQNTQRKWEAAGAAEQHRTYLEGECLEWLRRYLENGKETLQRA
AG1*05:01  ....I..............................................F........................................
AG2*01:01  ....I.............................I.........................................................
AG2*01:02  ....V.............................I.....................................K...................
AG3*02:01  ....I.......................................................................................
AG3*03:01  ....F.............................I.........................................................
AG4*08:01  ....L.............................V............L............R...............................
AG4*09:02  ....V.............................V............V............R...............................
AG5*07:06  ....V.............R........................................EDV..............S...............
AG5*07:10  ....I.............R........................................EDV..............S...............
AG6*04:01  ....V.........................S.............................R......................K........
E*02:01    ....L.W.H.........F...........K..LT........S.V.T...ISEQ.SNDGSE..HQ.A...DT.V...H............S
F*02:01    ....L.G.N...M..........H.H....K...S............TV.RI...FY..EEY..EF.I........L...............
I*01:01    ....L...C..............Y.S....................GE.............E...R.A....R.V........K........

α3 Domain        190     200       210       220       230       240           250       260       270
Consensus  DPPKTHVTHHPVSDYEATLRCWALGFYPAEITLTWQRDGEEQTQDTELVETRPTGDGTFQKWAAVVVPSGEEQRYTCHVQHEGLPKPLTLRW
AG1*05:01  ...........................................E...............................................F
AG2*01:01  ................................................M......N....................................
AG2*01:02  ......L....I..H.................................M......N....................................
AG3*02:01  .....N.....................................E.........................................E......
AG3*03:01  .....N.....................................E.........................................E......
AG4*08:01  .....................................................................................E......
AG4*09:02  .....Q......................................................................................
AG5*07:06  .....N.......................G..............................................................
AG5*07:10  ..............H..............G................................V.............................
AG6*04:01  ..............H............................E................................................
E*02:01    E.............H.................V.......D............A...............................E......
F*02:01    ....A..A...I..R......................................A...............................Q......
I*01:01    ..............H......................................G...N...........................E......
```

**B**

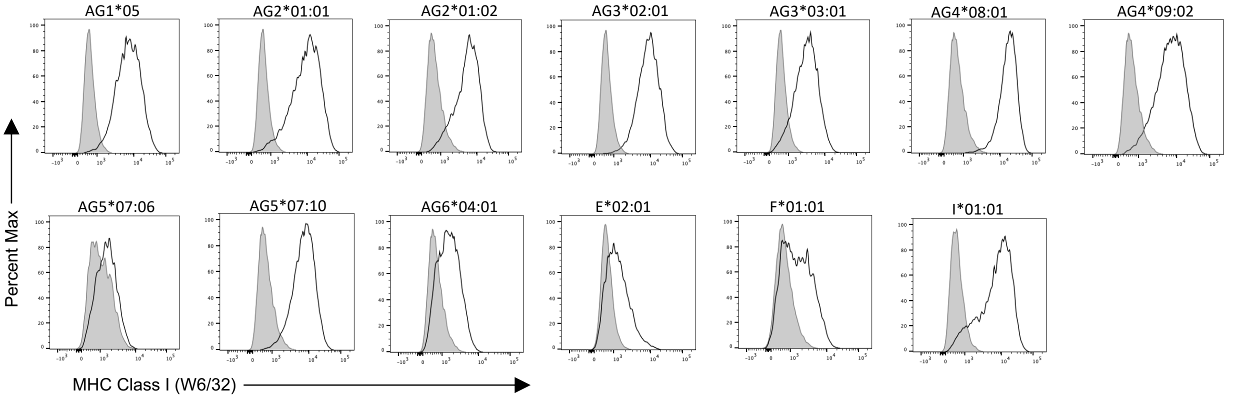


**Supplemental Figure 3.** (A) Amino acid alignment of the α1- α3 domains of non-classical Mamu molecules used in this study. Residues 77-83 corresponding to the Bw4/Bw6 motif are underlined and predicted KIR contact sites based on the crystal structure of HLA-B*57 in complex with KIR3DL1 are shaded (90). (B) Surface expression of MHC class I molecules on 721.221 cells. 721.221 cell lines expressing the indicated rhesus macaque MHC class I molecules (open) were stained with a PE-conjugated pan-MHC class I–specific mAb (clone W6/32) and compared with staining on parental 721.221 cell lines (shaded).

Supplemental Figure 4

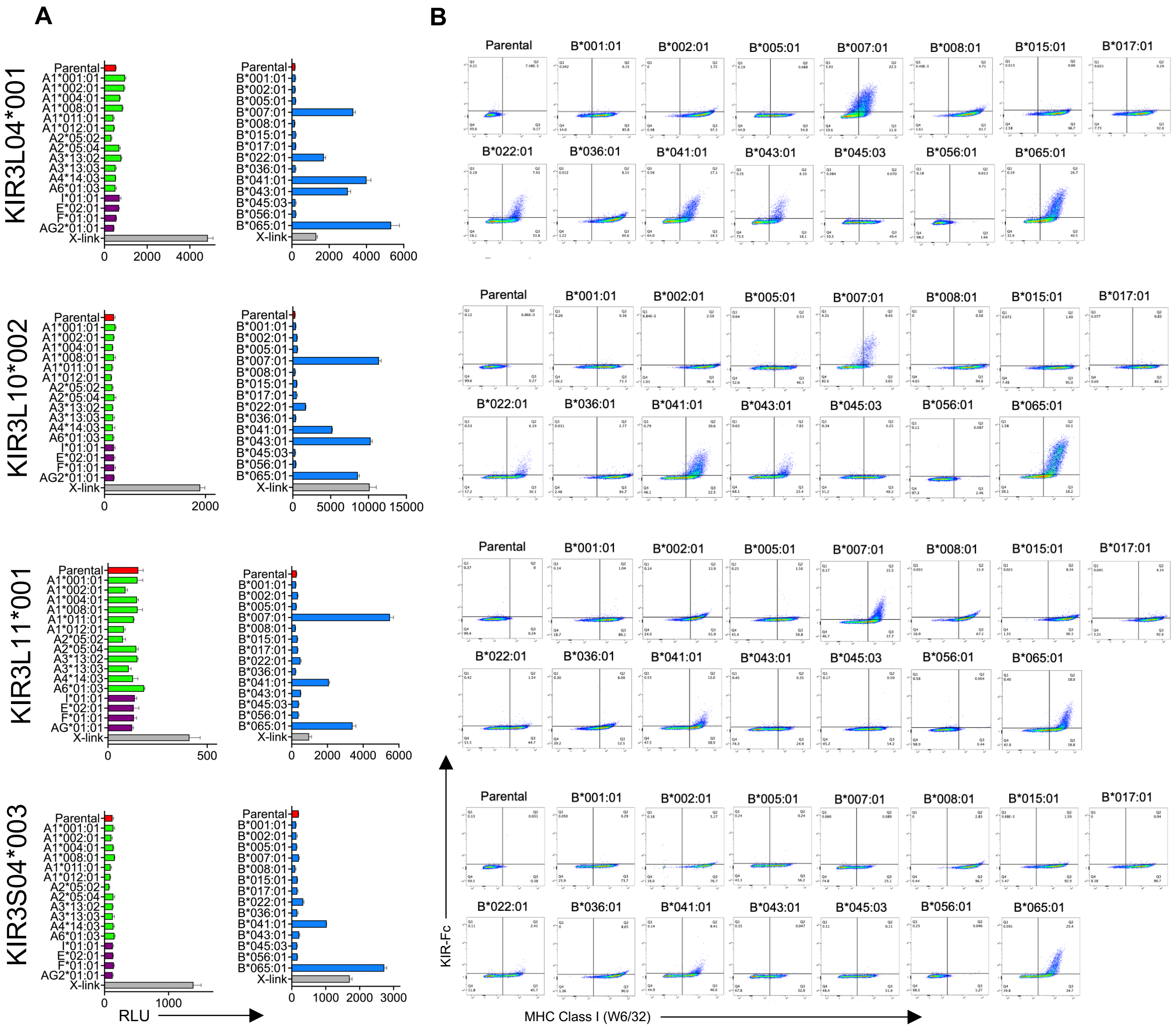


**Supplemental Figure 4.** KIR3D04*001, KIR3DL10*002, KIR3DL11*001 and KIR3DS04*003 interact with multiple Mamu-Bw4 ligands. (A) KIR3DL04*001-, KIR3DL10*002-, KIR3DL11*001- and KIR3DS04*003-CD3ζ JNL cells were incubated with 721.221 cells expressing the indicated rhesus macaque MHC class I molecules. Ligand recognition was detected by the MHC class I-dependent upregulation of luciferase (RLU). Bars are color coded according to products of the classical Mamu-A (green), -B (blue), and non-classical Mamu-I, -E, -F, and -AG (purple) genes. Parental 721.221 cells (red) and antibody crosslinking (X-link) were included as controls. Error bars indicate SD of the RLU values for triplicate wells. The data shown are representative of at least three independent experiments. (B) Parental 721.221 cells and Mamu-B+721.221 cells were stained with Near-IR LIVE/DEAD dye, KIR3DL04*001-, KIR3DL10*002-, KIR3DL11*001- and KIR3DS04*003-Fc followed by goat anti-mouse IgG and MHC class I-specific antibody (W6/32).

Supplemental Figure 5

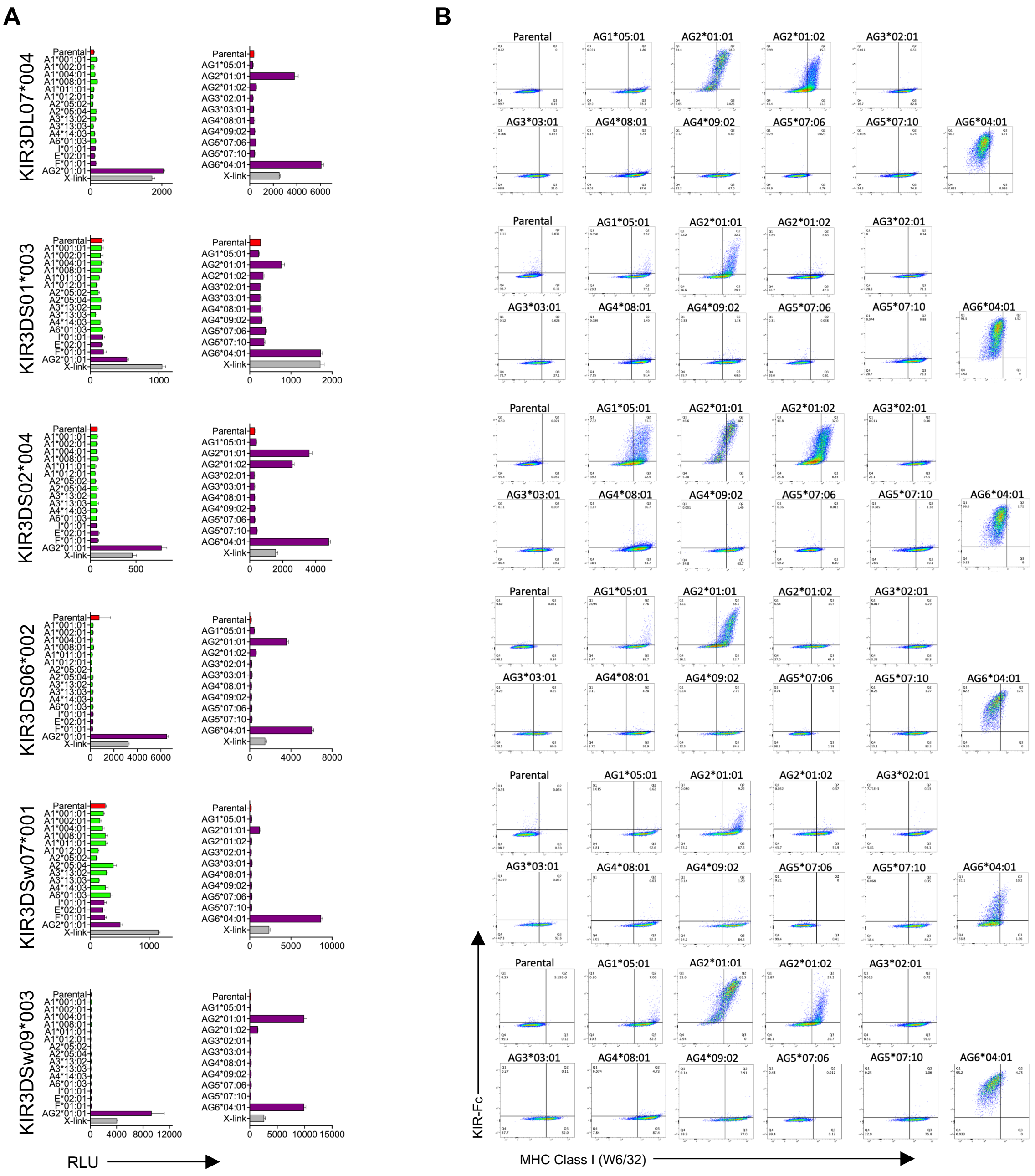

**Supplemental Figure 5.** KIR3DL07*004, KIR3DS01*003, KIR3DS02*004, KIR3DS06*002, KIR3DSw07*001 and KIR3DSw09*003 interact with Mamu-AG ligands. (A) KIR3DL07*004-, KIR3DS01*003-, KIR3DS02*004-, KIR3DS06*002-, KIR3DSw07*001- and KIR3DSw09*003-CD3ζ JNL cells were incubated with 721.221 cells expressing the indicated rhesus macaque MHC class I molecules. Ligand recognition was detected by the MHC class I-dependent upregulation of luciferase (RLU). Bars are color coded according to products of the classical Mamu-A (green) and non-classical Mamu-I, -E, -F, and -AG (purple) genes. Parental 721.221 cells (red) and antibody crosslinking (X-link) were included as controls. Error bars indicate SD of the RLU values for triplicate wells. The data shown are representative of at least three independent experiments. (B) Parental 721.221 cells and Mamu-B+721.221 cells were stained with Near-IR LIVE/DEAD dye, KIR3DL07*004-, KIR3DS01*003-, KIR3DS02*004-, KIR3DS06*002-, KIR3DSw07*001- and KIR3DSw09*003-Fc followed by goat anti-mouse IgG and MHC class I-specific antibody (W6/32).

Supplemental Figure 6

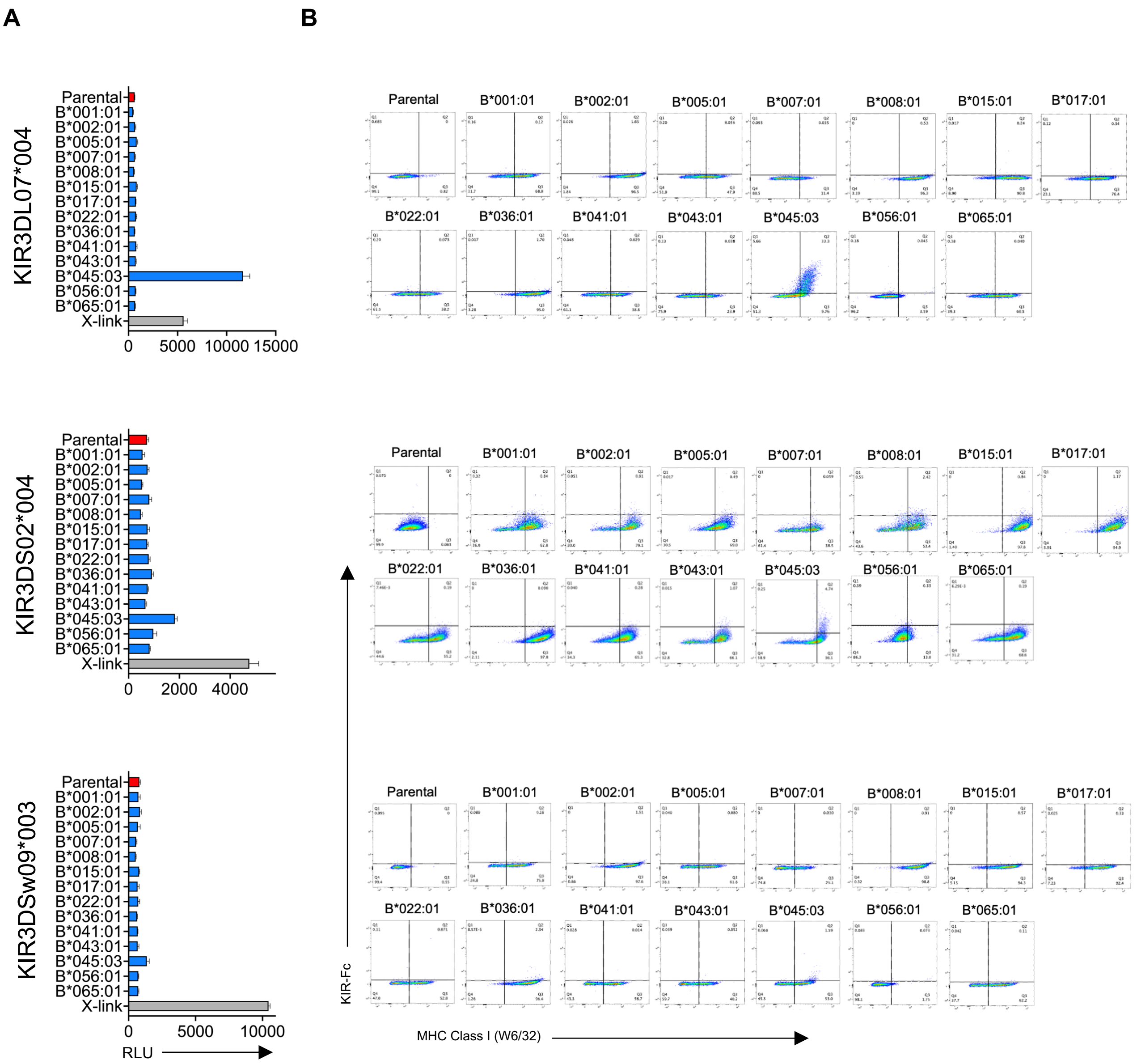


**Supplemental Figure 6.** KIR3DL07*004, KIR3DS02*004, and KIR3DSw09*003 interact with Mamu-B*045:03 (A) KIR3DL07*004-, KIR3DS02*004-, and KIR3DSw09*003-CD3ζ JNL cells were incubated with 721.221 cells expressing the indicated rhesus macaque MHC class I molecules. Ligand recognition was detected by the MHC class I-dependent upregulation of luciferase (RLU). Bars are color coded according to products of Mamu-B (blue) genes. Parental 721.221 cells (red) and antibody crosslinking (X-link) were included as controls. Error bars indicate SD of the RLU values for triplicate wells. The data shown are representative of at least three independent experiments. (B) Parental 721.221 cells and Mamu-B+721.221 cells were stained with Near-IR LIVE/DEAD dye, KIR3DL07*004-, KIR3DS02*004-, and KIR3DSw09*003-Fc followed by goat anti-mouse IgG and MHC class I-specific antibody (W6/32).

Supplemental Figure 7

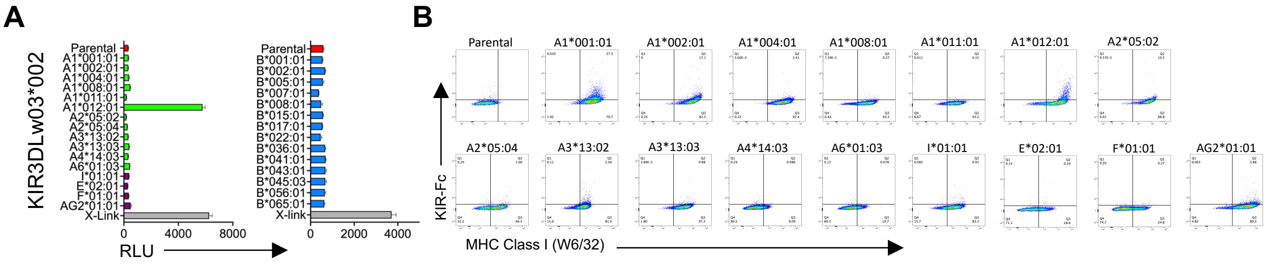


**Supplemental Figure 7.** KIR3DLw03*002 interacts with Mamu-A1*012:01. (A) KIR3DLw03*002-CD3ζ JNL cells were incubated with 721.221 cells expressing the indicated rhesus macaque MHC class I molecules. Ligand recognition was detected by the MHC class I dependent-upregulation of luciferase (RLU). Bars are color coded according to products of the classical Mamu-A (green), -B (blue), and non-classical Mamu-I, -E, -F, and -AG (purple) genes. Parental 721.221 cells (red) and antibody crosslinking (X-link) were included as controls. Error bars indicate SD of the RLU values for triplicate wells. The data shown are representative of at least three independent experiments. (B) Parental 721.221 cells and Mamu-B+721.221 cells were stained with Near-IR LIVE/DEAD dye, KIR3DLw03*002-Fc followed by goat anti-mouse IgG and MHC class I-specific antibody (W6/32).

Supplemental Figure 8

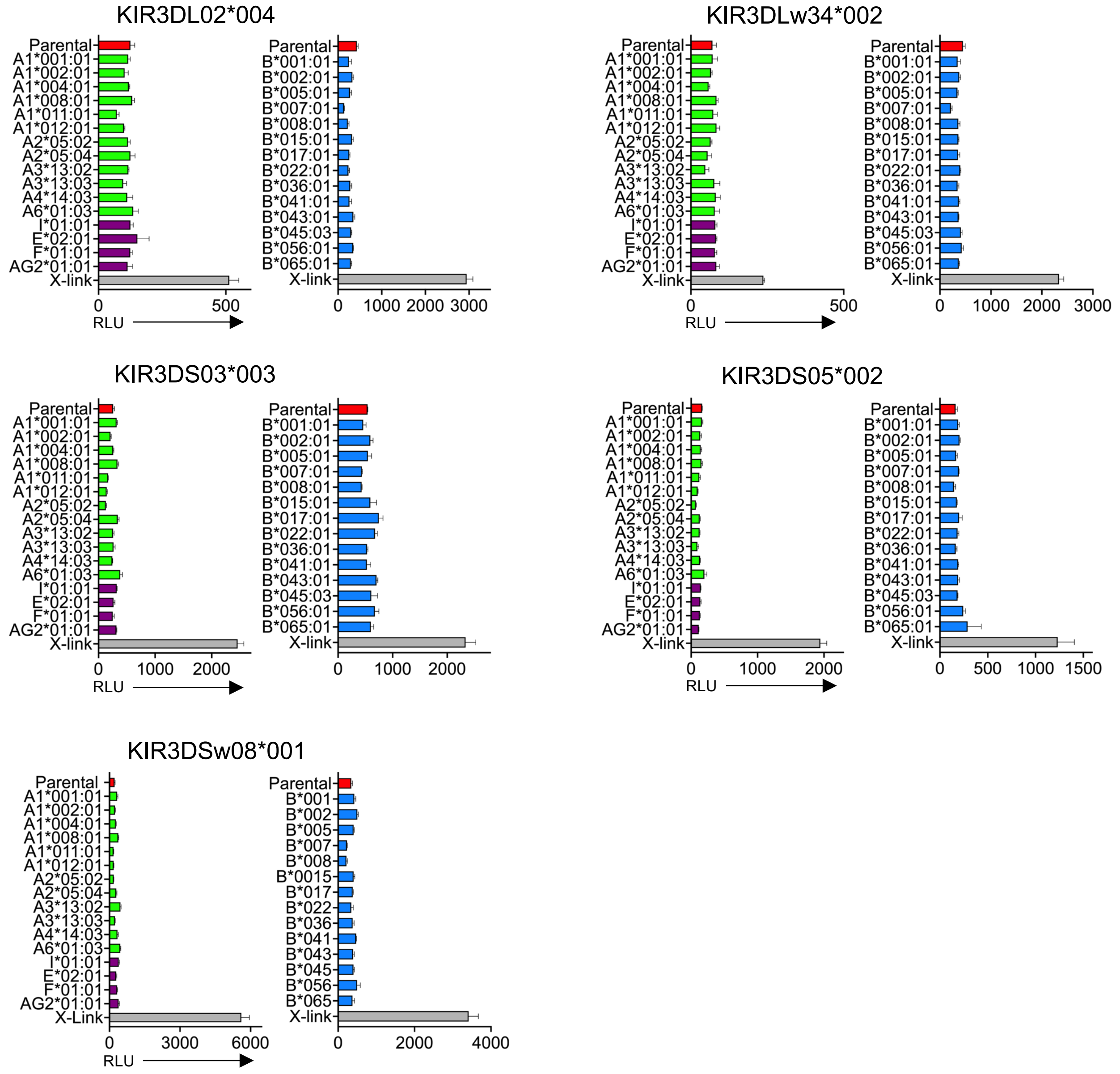


**Supplemental Figure 8.** Five of the KIR tested were found not to interact with any of the MHC class I ligands tested in this study. KIR3DL02*004-, KIR3DLw34*002-, KIR3DS03*003-, KIR3DS05*002-, and KIR3DSw08*001-CD3ζ JNL cells were incubated with 721.221 cells expressing the indicated rhesus macaque MHC class I molecules. Ligand recognition was detected by the MHC class I-dependent upregulation of luciferase (RLU). Bars are color coded according to products of the classical Mamu-A (green), -B (blue), and non-classical Mamu-I, -E, -F, and -AG (purple) genes. Parental 721.221 cells (red) and antibody crosslinking (X-link) were included as controls. Error bars indicate SD of the RLU values for triplicate wells. The data shown are representative of at least three independent experiments.
